# Supplemented nutrition increases immunity and drug efficacy in a natural host-helminth system

**DOI:** 10.1101/837617

**Authors:** Amy R. Sweeny, Melanie Clerc, Paulina A. Pontifes, Saudamini Venkatesan, Simon A. Babayan, Amy B. Pedersen

## Abstract

Gastrointestinal helminths are common parasites of humans, wildlife, and livestock, leading to chronic infections in large parts of the world. In humans, there is also an overlap in the incidence of malnutrition and helminth infections which can predispose individuals to higher infection burdens and reduced anthelmintic efficacy due to compromised immunity. This relationship has been well-studied in laboratory models by testing for the impact of dramatic reductions of specific nutrients on infection outcomes. However, much less is known about the benefits of whole-diet supplementation in natural host-helminth systems. We experimentally supplemented the diet of wood mice (*Apodemus sylvaticus)* and measured resistance to the gastrointestinal nematode *Heligmosomoides polygyrus* and anthelmintic treatment efficacy in both natural and captive populations. Supplemented wood mice were more resistant to *H. polygyrus* infection, cleared worms more efficiently after treatment, produced stronger general and parasite-specific antibody responses, and maintained better body condition. In addition, supplemented nutrition in conjunction with anthelminthic treatment significantly reduced *H. polygyrus* transmission potential. These large-scale improvements in condition and immunity of supplemented nutrition found in controlled and wild environments show the rapid and extensive benefits of a well-balanced diet and have important implications for using diet interventions to improve disease control programmes.

**Author Summary:** Gastrointestinal helminths are ubiquitous parasites which cause chronic infections and debilitating symptoms worldwide for human and animal populations alike. Efforts to control helminth infections rely primarily on deworming drugs. However, despite high availability and low cost of these drugs, reinfection is common and helminths remain a substantial problem in many areas of the world. One factor which contributes to the persistence of helminth infections is co-occurrence of malnutrition. Extensive work in model laboratory systems has shown that deficiencies in both macro- and micronutrients can impair host immune responses to helminth infections, but how this relationship translates to natural host-helminth systems and drug efficacy is still poorly understood. Here, we used a combination of experimental nutrition supplementation and deworming treatment in paired wood mouse populations in both wild and captive environments to determine the role of a well-balanced diet for infection control in a natural host-helminth system. We found that mice on a supplemented diet had lower burden of infection and increased drug efficacy, body condition, and immune responses. We also found that supplementation lowered transmission potential of hosts. These broad and rapid effects of increased nutrition quality have important implications for the benefits of diet interventions alongside deworming drugs.

## Introduction

Gastrointestinal (GI) helminth parasite infections are ubiquitous and one of the most common causes of chronic disease in wildlife, livestock, and human populations [1]. Globally, helminths infect one in three humans [2], often causing stunted growth and development, impaired physical fitness and cognition in children, pregnancy complications, and decreased productivity in livestock [1–5]. Among wildlife, helminth infections are very common and can significantly impact host survival and reproduction, while also playing a key role in regulating population cycles and dynamics [6–8]. To reduce the burden (number of worms per individual) and morbidity of helminth infections, standard treatment in humans and livestock is drug therapy [9, 10]. However, despite the high availability and low cost of anthelmintic drugs, morbidity from helminth infections remains high [2]. Further, reinfection post-treatment is rapid [11].

The most common human helminth infections are caused by gastrointestinal nematodes [2], whose transmissible stages can persist for long periods of time in the environment, especially in areas with poor sanitation [12]. For example, within one year of treatment in human populations where worms are endemic, *Ascaris lumbricoides* prevalence can reach nearly 100% of pre-treatment levels, while *Trichuris trichiura* and hookworm can reach 50% of pre-treatment prevalence [13, 14]. Effective helminth control therefore depends not only on reducing burdens within all individuals, but also reducing exposure and susceptibility to reinfection. To achieve this, a better understanding of both the environmental and host factors that drive reinfection is needed [15].

Limiting environmental exposure by integration of sanitation and hygiene improvement [16] or grazing management in livestock [17, 18] can be an effective strategy for nematode control, but these methods do not address underlying susceptibility to infection or treatment efficacy. Individual differences in response to infection and treatment are largely mediated by immune and nutritional status [19, 20]. It is well-established that micronutrient, macronutrient, and overall energy deficiencies impair the immune system [21] and given that mounting an immune response is costly, insufficient resources can worsen the impact of nematode infection [22]. Additionally, pre-existing malnutrition can also increase susceptibility to infection and/or compromise immune responses [20]. These relationships have largely been studied in humans in areas with endemic nematode infections and malnutrition [19, 23], but also exist in other organisms. In livestock, the increased nutrition demand of late pregnancy and lactation is associated with a substantial increase in GI nematode burdens [24] and in wild animal where resource availability is typically limited, trade-offs between the immune response and other energetically costly processes mean that individuals must divert resources between different physiological demands [22].

Integration of nutritional supplementation with drug therapies has been explored to address the problem of reinfection after treatment [25, 26], and clinical trials in humans have measured the impact of single or combined micronutrient supplements [27–31] on the reduction of *Ascaris, Trichuris,* and hookworm burdens. However, a recent meta-analysis found contradictory results, reporting both positive and negative impacts of supplemented nutrition on nematode infection [32]. Laboratory mouse models have shown that both macro- and micro-nutrients play a key role in host immunity to nematodes and susceptibility to infection [33–37]. For example, deficiencies in protein [35] and zinc [33, 34] have been shown to increase worm burdens and reduce eosinophilia and parasite-specific IgG1 response [33] to *Heligmosomoides bakeri* (formerly *Heligmosomoides polygyrus bakeri*), a well-studied model nematode. However, laboratory conditions are highly controlled and therefore unlikely to mimic life in the wild while laboratory house mice (*Mus musculus domesticus*) do not have a long co-evolutionary history with *H. bakeri* and thus may not reflect natural infection dynamics [38]. Although effects of experimental supplementation have been investigated in wild mouse models [7,39,40], these studies supplemented the food supply to mimic natural variation rather than increased food quality.

Here we experimentally enriched nutrition, by supplementing mice with a well-balanced diet, to test the impacts on immunity to *H. polygyrus* and anthelmintic treatment efficacy in wood mice (*Apodemus sylvaticus*). Wood mice live in woodlands across much of Europe and are commonly, chronically infected with *H. polygyrus* (prevalence 20-100%) [41, 42], a sister taxa to *H. bakeri* [43]. Importantly, while anthelmintics can significantly reduce infections in wood mice, reinfection to pre-treatment burdens occurs within 2-3 weeks [44, 45]. Further, wild wood mice have significant energetic demands for reproduction, foraging, and survival, and are exposed to marked seasonal changes [46, 47]; conditions which laboratory settings cannot replicate, but which are likely drivers of infection exposure, immunity, and nutrition availability. Crucially, here we have the unique ability to test same host-helminth system in both the wild and controlled laboratory conditions in order to control infection/reinfection, exposure, coinfection and other important factors, as we have a wild-derived, but now laboratory reared, colony of wood mice and a wild-collected *H. polygyrus* isolate [48]. We experimentally tested the effects of supplemented nutrition and anthelmintic treatment on (i) *H. polygyrus* burden and egg shedding in the wild (ii) egg shedding, susceptibility to infection and reinfection in the laboratory and (iii) body condition and immune responses in the wild and in the same species in controlled laboratory settings. We found strong evidence of rapid and broad impacts of this well-balanced diet for host condition, helminth resistance and treatment efficacy; suggesting that whole-diet supplementation could provide significant benefits for helminth control by increasing the host’s ability to respond to infection and reducing the probability of reinfection.

## Materials and Methods

### Ethics Statement

All animal work was conducted in accordance to UK Home Office in compliance with the Animals (Scientific Procedures) Act 1986, was approved by the University of Edinburgh Ethical Review Committee, and was carried out under the approved UK Home Office Project License 70/8543. Animal sacrifice was performed using Supplemented nutrition increases immunity against helminths Schedule 1 Methods (Field: Cervical dislocation; Laboratory: Rising CO2 exposure). Fieldwork was carried out with permission of the Forestry Commission Scotland. Permit reference SUR09.

### Field experiment

#### Experimental Design

We conducted the field experiment in a 100ha broadleaf woodland in Falkirk, Scotland (Callendar Wood, 55.990470, −3.766636). This woodland has wood mouse (*A. sylvaticus*) populations that are naturally infected with *H. polygyrus* [45]. We conducted two 8-week experiments (2 temporal replicates) during the wood mouse breeding season, when host energetic demands are highest: (i) May - July 2015 and (ii) June - August 2016. We used a 2 x 2 factorial design, where nutrition was manipulated at the population level (unit: trapping grid; see SI): supplemented high-quality, whole-diet food pellets (hereafter “diet”) vs. control (unmanipulated), while anthelmintic treatment (hereafter “treatment”) was manipulated at the individual level (unit: mouse; control (water) vs. treatment; Fig. 1). We live-trapped mice for 3 nights/week using Sherman live traps (H.B. Sherman 2×2.5×6.5-inch folding trap, Tallahassee, FL, USA). All wood mice weighing >10g at first capture were tagged with a subcutaneous microchip transponder for identification (Friend Chip, AVID2028, Norco, CA, USA). On both control and diet-supplemented grids, all mice at first capture were rotationally assigned, within each sex, to either control or drug treatment groups.

**Figure 1.**
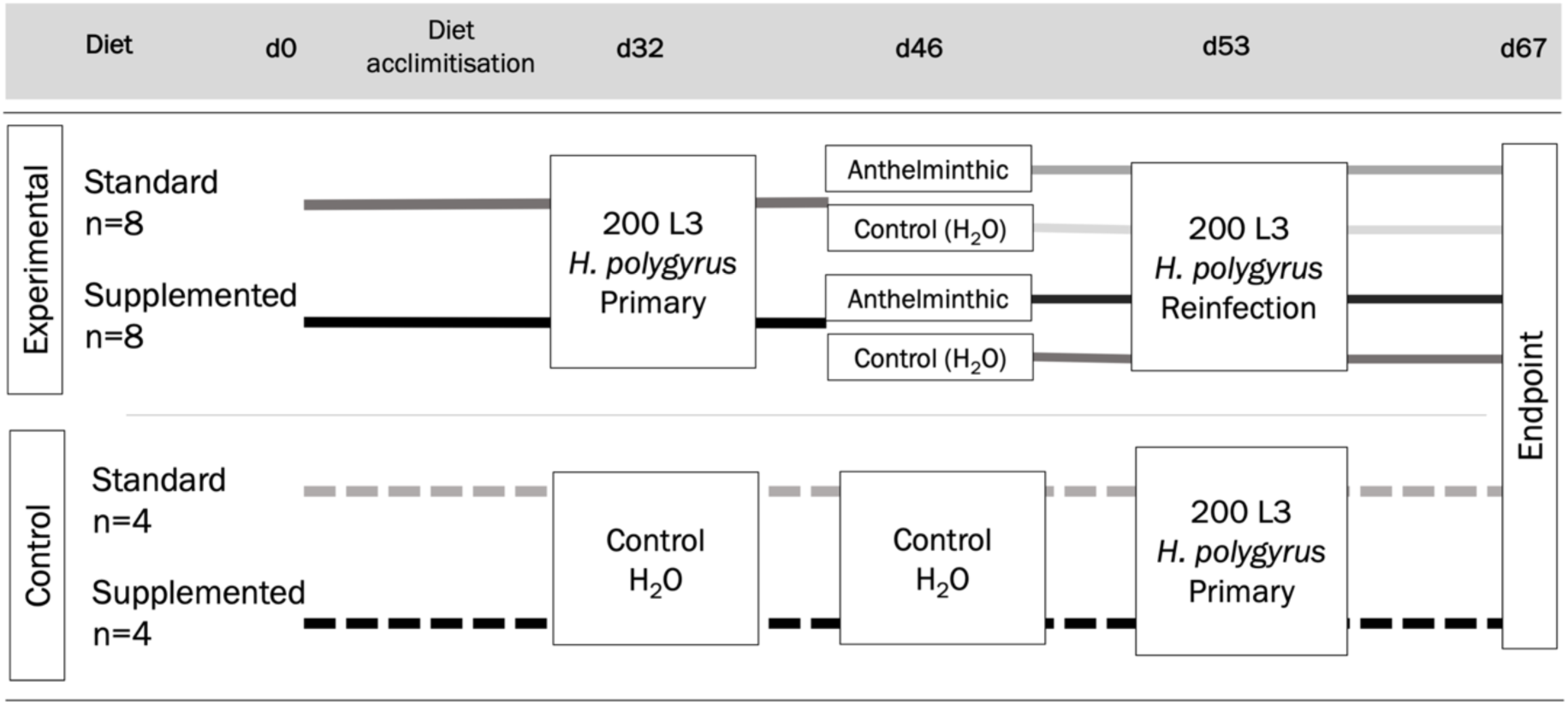
Diagram of the laboratory experimental design. All wood mice were assigned to diet groups at day 0 (d0). After 32 days of diet acclimatisation (d32), all experimental (n=16; solid lines) mice were given a primary challenge of *H. polygyrus*. On day 46, 14 days post infection (dpi), the experimental mice were randomly assigned within diet groups to either receive an anthelminthic treatment (darker lines) or a control dose of water (lighter lines; n=4/ group). On day 53, all experimental mice received a secondary challenge of *H. polygyrus*, and were culled on day 67, 14 days post-secondary challenge. On days 32 and 46, control mice (dashed lines, n=4/ group) received equivalent volumes of water, and on day 53 received a primary challenge with *H. polygyrus*. Control mice were culled on day 67, 14 days post primary challenge.

#### Food supplementation

We diet-supplemented for two weeks before and then throughout the 8-week experiment twice per week with 2kg/ 1000m^2^ of sterilised, TransBreed™ mouse chow pellets, scattered at regular intervals across the grids to ensure an even spatial distribution. TransBreed™ is a high-nutrient, standard veterinary feed which is formulated for optimum breeding performance in laboratory mice and offers whole-diet nutrition to the wild mice in this study (20% protein, 10% fat, 38% starch, high content of micronutrients, full details in Table S1). Our supplementation complemented natural food availability.

#### Anthelminthic treatment

We administered a weight-adjusted (2ml/g) dose of Pyrantel pamoate (Strongid-P, 100 mg/kg) and Ivermectin (Eqvalan, 9.4mg/kg) to each mouse allocated to the anthelminthic group, a combination dose shown to be highly effective at removing both adult and larval *H. polygyrus* from wood mouse for 12-16 days in our previous work [45].

#### Sampling

For each mouse at every capture we measured: sex, age (determined by size and coat colour, as described in SI), and host condition including body mass, length, fat scores, and reproductive status (as described in SI). Blood samples were collected via mandibular bleed (first capture), tail snip (subsequent captures), or terminal bleed (sacrificed animals) a maximum of once per week. Faecal samples were collected for each mouse at every capture from previously sterilised traps. Mice were sacrificed 12-16 days after first capture, corresponding to the period of efficacy for these drugs in wild wood mice [52, 55]. Mice caught outside of this date range, or those pregnant or lactating at capture, were released. Eyes were collected as a proxy of age (details in SI) and small intestine, caecum, and colon of each individual were collected for counts of adult *H. polygyrus* worms.

### Laboratory experiment

#### Experimental design

We conducted a 2×2 factorial design in a laboratory colony of *A. sylvaticus* (details in SI), parallel to our field experiment: (i) diet (standard vs. supplemented, both Special Diet Services (SDS) pelleted rodent chow) and (ii) anthelmintic treatment (control (water) vs. treatment); both implemented at the individual level (Figure 1). As in the field experiment, Transbreed™ was used for the diet supplementation. Rat Mouse 1 (RM1™) was used as the control diet, as it is a maintenance chow with lower nutrient content, but is not considered a restrictive diet (Table S1). Mice on both diets were fed *ad libitum* and were given a 32-day diet acclimatisation period. Sixteen wood mice from our colony, aged 15-21 weeks (median 18 weeks), were randomly assigned to 4 experimental groups (n = 4/group; Figure 1): (i) supplemented nutrition, treated, (ii) supplemented nutrition, control, (iii) standard nutrition, treated, and (iv) standard nutrition, control. Eight mice were designated as controls and placed on the same standard and supplemented mice as experimental mice (n = 4 per group; Figure 1).

#### Infections and anthelminthic treatments

We performed primary and secondary *H. polygyrus* infections to mimic the high level of exposure found in wild wood mice. After diet acclimatisation, all experimental mice were infected with 200 wild-derived *H. polygyrus* L3 in 150uL dH_2_O via oral gavage. On day 46, 14 days post-infection, half of the male and female mice challenged with *H. polygyrus* were randomly assigned to treatment groups and were given either anthelmintic drug treatment (identical combinations and doses as in field experiment, details in SI) or a control dose of water via oral gavage. Control animals received equivalent water control on day 32 and 46 experimental primary infection and treatment timepoints. On day 53, all experimental mice were re-infected with 200 *H. polygyrus* L3 in 150uL H_2_O via oral gavage to act as a secondary challenge, and control mice were given a primary challenge with 200 *H. polygyrus* L3. On day 67 (14 days post-secondary challenge) all mice were culled and adult *H. polygyrus* in the small intestine were counted.

#### Sampling

Once per week, both weight (grams) and fat scores (described in SI) were recorded. From the day of primary infection (day 32) we collected a weekly blood sample for each individual; on days 32, 39, and 53 the samples were taken via venesection (tail bleed), while on day 46, the samples were taken via venepuncture (cheek bleed). On day 67, individuals were sacrificed and sampled as described above. Faecal samples were collected three times/week starting at primary infection and then continued throughout the course of the experiment. Over the course of this experiment, 5 mice exhibited weight loss over the threshold for our experimental protocol, not related to the diet supplementation or *H. polygyrus* infection and were culled and removed from further analysis.

### Laboratory assays for both field and laboratory experiments

*H. polygyrus* egg shedding was measured as eggs per gram of faeces (EPG). using salt flotation and microscopy following Knowles et al. (2013) (details in SI). We used ELISA assays to measure (1) total faecal IgA concentration and (2) sera *H. polygyrus-*specific IgG1 antibody titres for each mouse at each capture/sampling point as previously described for wood mice [56] (see SI for details). We calculated total faecal IgA concentration by extrapolation from a standard curve of known concentrations from a synthetically manufactured standard antibody. *H. polygyrus-*specific IgG1 was calculated as a relative concentration to a positive reference sample as described in the SI. We refer to both IgA and IgG1 values as ‘antibody concentration’.

### Statistical Analysis

We carried out all statistical analysis using R v 3.5.1 [49]. All models were fit using the package ‘glmmTMB’ and are listed in Table S2. Post-hoc comparisons of interaction levels were calculated using the package ‘emmeans.’

#### *H. polygyrus* infection

To investigate the impact of supplemented nutrition on *H. polygyrus*, we used Generalized Linear Models (GLMs) for the following response variables: (i) intensity of infection (EPG) at first capture (before treatment) (ii) mean EPG per individual for subsequent post-treatment captures and (iii) infection burden (adult worm count) at final capture. The distribution of EPG intensity and worm burden were highly over-dispersed, which is typical for many helminth infections [8], thus we fit the models with negative binomial (NB) error distributions. Fixed effects in all models included diet and host characteristic variables (i.e. sex, reproductive status, age full details in Table S2). Drug treatment and a treatment - diet interaction were included in models of data after first capture. Age was only included as an explanatory variable in the model examining worm burden, where lethal sampling allowed the use of eye lens weight as a proxy for age [57]. Diet were classified as ‘supplemented’ if mice were captured > 50% of the time on supplemented grids and as ‘control’ otherwise. Only 18% (n = 16) of mice were captured on both grid types, but to test the possibility that effects of supplemented nutrition could be dependent on the time spent on these grids, we fit another set of 3 models, identical to those described above, but including three levels of diet as an explanatory variable (control, mix, supplemented), where ‘mix’ represented sixteen mice that were found on both control and supplement grids across the experiment (see SI).

We investigated the impact of supplemented nutrition and treatment on *H. polygyrus* infection in the laboratory using GLMs with NB errors to the following response variables to primary infection: (i) peak EPG shed, (ii) total EPG shed, and (iii) adult worm burden. Although all experimental mice experienced primary and secondary infection, only two individuals had EPG values > 0 after reinfection, and models could not be reliably fit to this dataset. Therefore, only adult worm burden was used as a response variable for secondary infection. Explanatory variables for all models are listed in Table S2, and included diet, host characteristics, and day of experiment as fixed effects. Although anthelmintic drugs were administered to half of the experimental group before secondary challenge, there was no difference in worm clearance (as indicated by EPG) between drug-treated and control mice (Figure S8B) and thus they were combined for these analyses.

#### Body condition

We fit GLMMs to two metrics of body condition to determine the effect of supplemented nutrition in the wild (i) body condition index (BCI, weight vs. length regression residuals; [50]) and (ii) total fat score (FS, sum of dorsal and pelvic fat scores [51]), both of which were normally distributed. Diet and drug treatment, host characteristics, and time were included as fixed effects, as well as *H. polygyrus* infection intensity (log of egg/gram+1; Table S2). We also fit a supplement-by-day interaction to investigate differences in the slope of the body condition-diet relationship over the 2-week period. Due to variation in weight change due to growth and gestation, we excluded non-adults and pregnant females from base body condition models, and fit separate models for pregnant females. Explanatory variables included in models for pregnant females were as above, except that sex and reproductive condition were excluded.

We fit linear mixed models (LMM) to determine the effect of supplemented nutrition on body condition in the laboratory. Because we expected less variable body length in the laboratory than in the wild, we did not take regular length measurements, therefore we only tested body mass (g) (not weight/ length residuals) and total fat score (as measured in the wild, details in SI) as response variables. We selected fixed effects to investigate the same relationships as in the wild (Table S2).

#### General and specific antibody measures

We tested for the effect of supplemented nutrition and *H. polygyrus* infection on general and specific antibody response in the wild and laboratory, by fitting GLMMs with either total non-specific IgA or *H. polygyrus-*specific IgG1 (for standardised IgG1 values >0 only) as the response variable with gaussian error distributions. Fixed effects included host characteristics, year and day of experiment, and experimental manipulations (Table S2) and all mixed models included mouse ID and grid:year as a random effect.

## Results

Our field experiment included 310 captures of 91 individual mice (2015: n = 49 and 2016: n = 42); 61 of which were captured > 1 time (mean number of captures = 3.42 +/− 0.26). Of these, 36 mice were sacrificed after two weeks to measure *H. polygyrus* adult worm burdens and for all other captures *H. polygyrus* eggs/gram was used as a proxy of adult worm burden (R_2015_ = 0.72, R_2016_ = 0.80, Fig S3). Adult mice comprised 87.5% of all captures in our dataset (juveniles = 2.4%, subadults = 10.1%). At the start of field and colony experiments, wood mice in the colony compared to wild wood mice had higher body mass (Colony mean weight = 23.88g; Wild mean weight = 20.32g; T-test, t = −2.99 p = 0.005) and better body condition (Colony mean total fat score = 9.08/10; Wild mean total fat score = 5.7/10, Wilcoxon Rank-Sum test, W = 127, p < 0.001).

### Supplemented nutrition decreased *H. polygyrus* worm burdens and egg shedding and improved anthelmintic drug efficacy

#### Field experiment

On average, mice spent ∼30 days (SEM +/− 1.5 days) on supplemented grids, but the range was between 12– 63 days. We found that mice caught for the first time on supplemented grids had significantly lower *H. polygyrus* EPG than mice on control grids (Figure 2A and 3A, β = −1.42, p = 0.018); resulting in ∼50% less egg shedding (23.55 vs 43.68 EPG, Table S4). By 12–16 days after first capture, mice on supplemented grids also had 60% fewer adult worms compared to mice on control grids (Figures 2C and 3A, β = −1.2, p = 0.045; Table S4).

**Figure 2.**
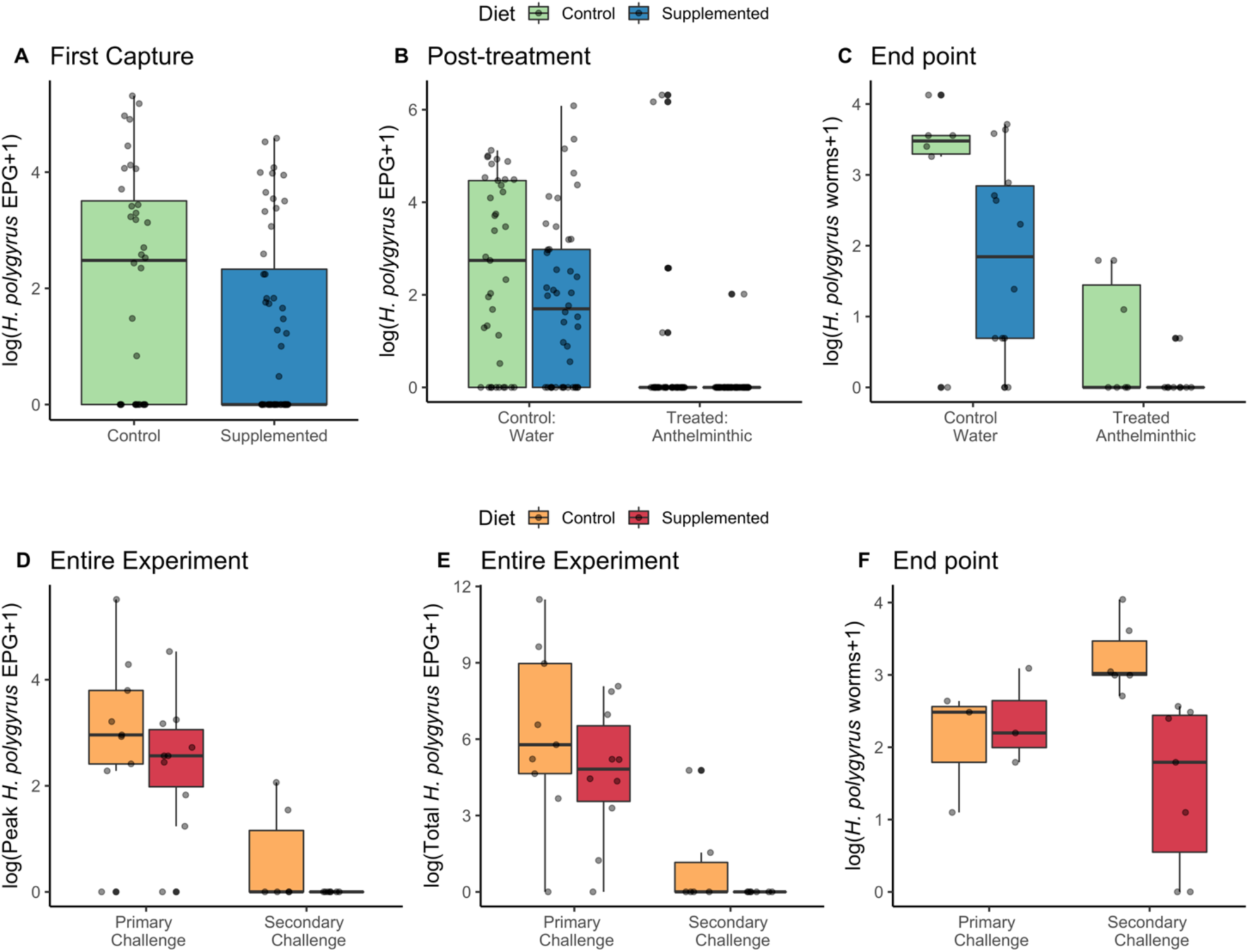
Effect of a supplemented nutrition diet on *H. polygyrus* infection in wild and laboratory wood mice. Mice on supplemented grids compared to control grids had lower mean *H. polygyrus* abundance (log EPG + 1) **(A)** at first capture, N = 88 individuals and **(B)** after treatment (N = 62 individuals; 166 captures) and had lower burden (log of worm count) **(C)** at end point for culled individuals (N = 36). Mice in the laboratory on a supplemented versus standard (control) diet had lower **(D)** peak shedding and **(E)** total shedding *H. polygyrus* EPG during primary and secondary challenge and had no difference in **(F)** worm burden (log of worm count) 14 days primary infection, but reduced worm burden 14 days post-secondary challenge with 200L3 *H. polygyrus.* N = 19 mice (Primary only control group, N = 6, Primary +Secondary group, N = 13). Black points represent outliers of boxplots; grey points represent raw data points.

**Figure 3.**
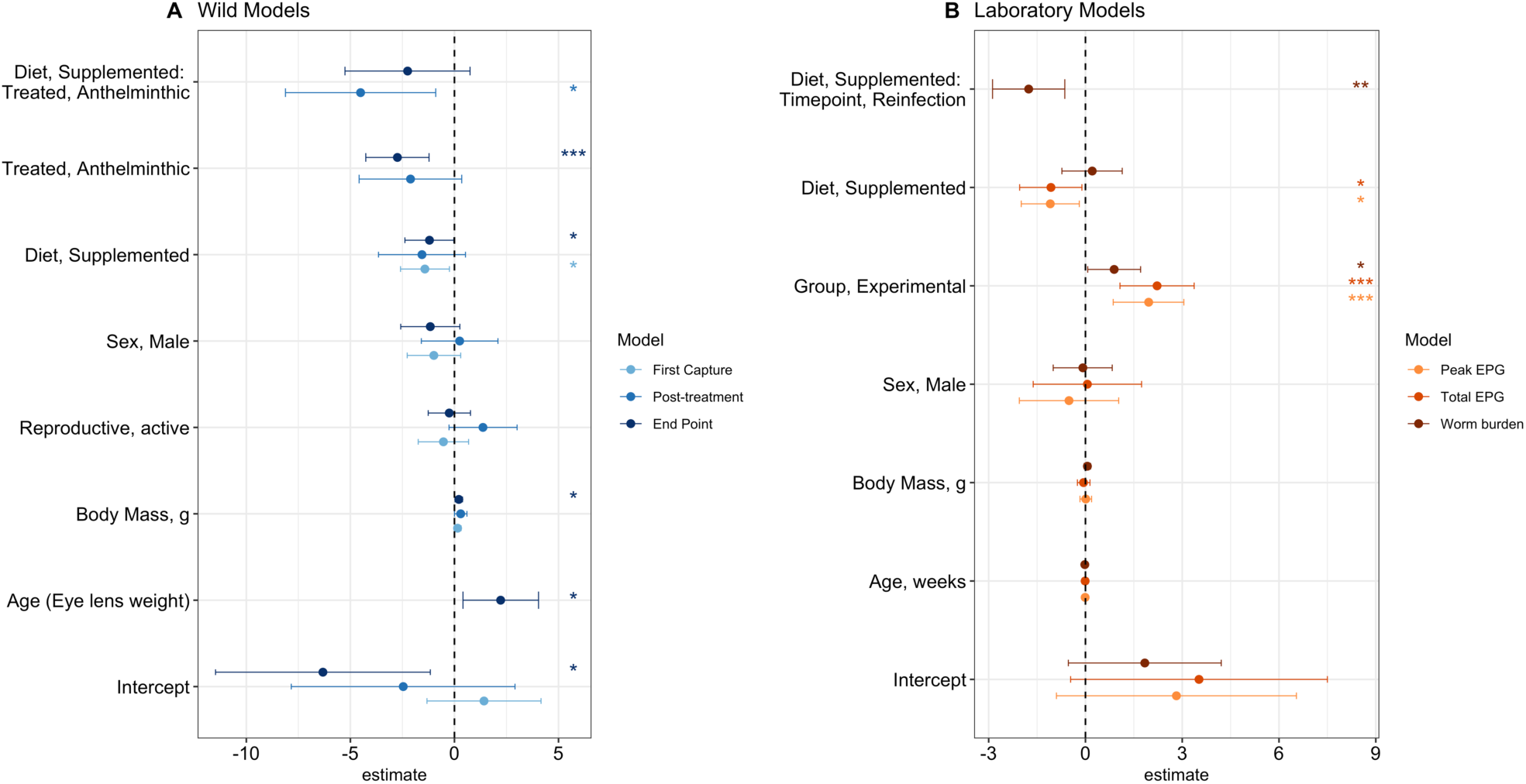
Effect size estimates from models investigating the effect of supplemented nutrition on *H. polygyrus* infection in (A) wild and (B) colony wood mice. Response variable and data used in each model is indicated by colour. In the wild, models represent faecal egg counts (EPG) for (i) first capture (light blue, GLM) and (ii) mean post-treatment captures (blue; GLM) and adult worm burdens for (iii) the experimental end point for culled animals (∼ 12-16 days post treatment; dark blue; GLM). In the laboratory, models represent (i) peak EPG (light orange, GLM), (ii) total EPG (orange, GLM), and adult worm burdens (dark orange, GLM). Diet*timepoint interaction was included in the endpoint models to investigate potential differences in effects of supplemented diet on primary and reinfection. Points and ranges represent model estimates and 95% credibility estimates for each model. Asterisks indicate the significance of variables: ***, ** and * indicate p < 0.001, p < 0.01 and p < 0.05 respectively. Eye lens mass (log-transformed) was included as a proxy for age in final capture model only.

In addition to main effects of diet supplementation alone, we also found a significant interaction between diet and anthelminthic treatment following assignment to treatment groups at first capture (EPG; β = −4.51, p = 0.01, Figure 3A). Notably, treatment reduced shedding to < 1 *H. polygyrus* egg per gram faeces in diet-supplemented mice for two weeks following treatment, while treated mice on control grids still shed ∼29 eggs/gram faeces during the same period compared to control grids (Tukey post-hoc test: β = −6.06, p < 0.001, Figure 2B). Likewise, for adult worms, although anthelmintic treatment significantly reduced worm burden for all mice (β = −2.74, p < 0.001), efficacy was highest in mice on supplemented grids (Tukey post-hoc test: β = −3.46, p = 0.017, Figures 2C and Figure 3A); resulting in complete worm clearance for all but one mouse, who had a single worm (Table S4).

For all models of *H. polygyrus* EPG in the wild, diet and a diet-by-treatment interaction were the only significant predictors (Figure 3A). For *H. polygyrus* worm burden, we found body mass and age to be additional predictors of infection, where larger (β = 0.21, p = 0.019) and older (β = 2.22, p = 0.016) individuals carried higher worm burdens (Figure 3A). Including a factor level accounting for a mixed amount of supplemented nutrition did not significantly improve the fit for models of EPG at first capture, post-treatment, or at end point (ΔAIC = 1.62, ΔAIC = −1.15, ΔAIC = 1.28, respectively; Fig S1) nor did the inclusion of this factor level change any of the main results (See supplementary information and supplementary figures S1 and S4)

**Figure 4.**
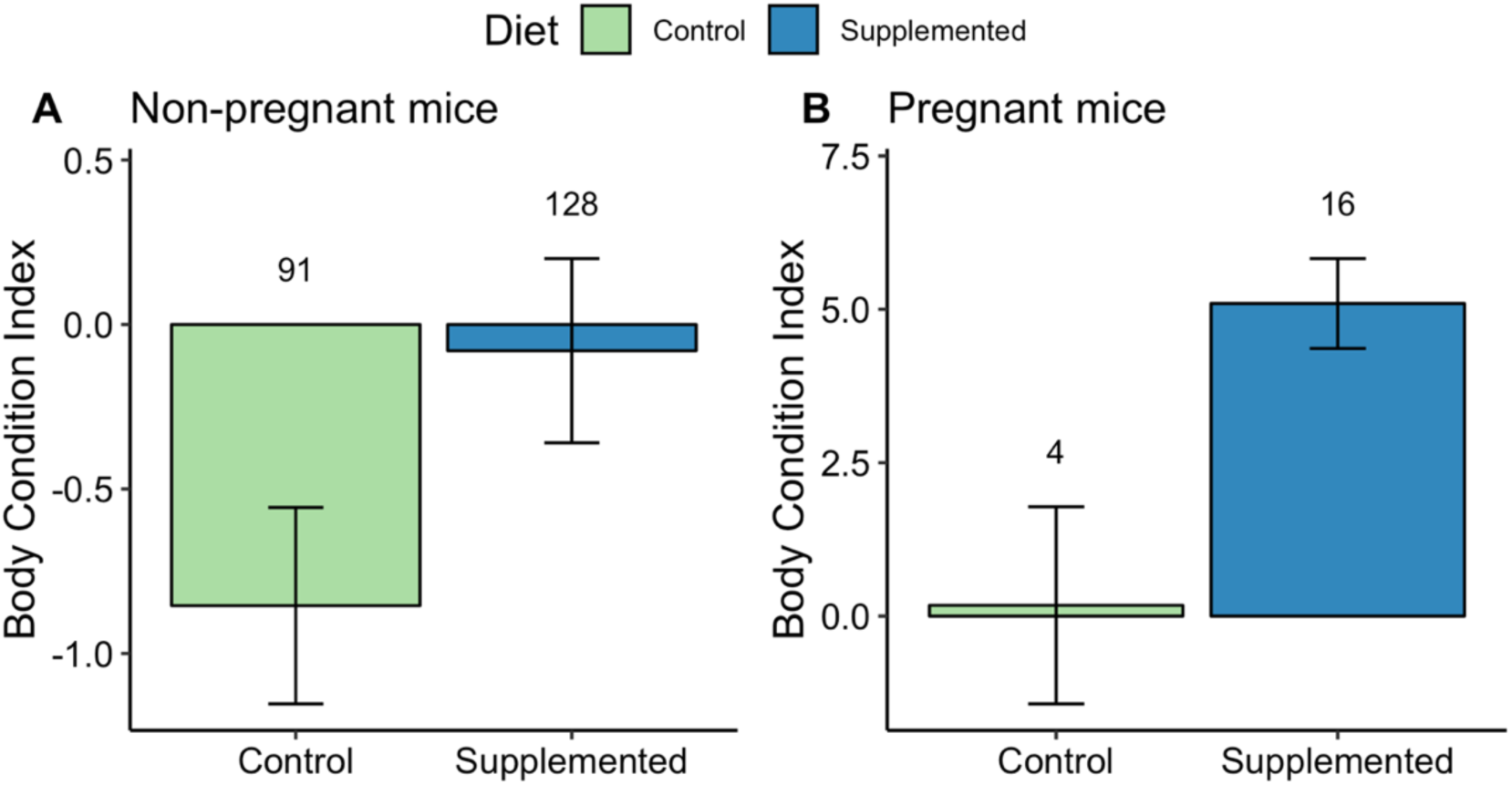
Effect of supplemented diet on body condition. Figure represents raw means +/− SEM for body condition index (weight/ length regression residuals). Numbers above bars indicate number of observations per group. Body condition index was higher in supplemented individuals for both A. Non-gestating mice B. Female mice gestating at the time of body condition measurement.

#### Laboratory experiment

In the controlled laboratory experiment, supplemented diet reduced both peak (β = −1.96, p = 0.017) and total *H. polygyrus* EPG (β = −1.07, p < 0.001) compared to mice on the standard ‘control’ diet, and, importantly mice receiving the diet supplement shed no eggs during reinfection (Figures 2D-E and 3B; Table S4). While there was no difference in adult worm burdens between mice on supplemented or control diets after primary infection, mice on the supplemented diet were significantly less susceptible to secondary challenge, with 75% lower adult worm burdens (β = −1.76, p = 0.002, Figures 2F and 3B; Table S4).

### Supplemented nutrition increased wood mouse body condition

Wild wood mice on supplemented grids had higher body condition (BCI) and total fat scores (FS) compared to mice on control grids (BCI β = 1.87, p = 0.003; FS β = 0.73, p = 0.005; Figures 4, S5); however, we also found significant interactions between diet and day of experiment, suggesting that supplemented mice lost weight faster than control mice over time despite overall greater condition (BCI: diet, supplemented*day β = −0.18, p = 0.009; FS: diet, supplemented*day β = −0.11, p = 0.002; Figure S5). In addition, we found that reproductively active mice had higher BCI scores (β = 1.24, p = 0.005); however, the relationship with FS was weakly negatively (β = −0.4, p = 0.052) (Figure S5). Among pregnant females, supplemented diet was also associated with significantly higher BCI and total FS (BCI β = 8.32, p < 0.001, FS β = 1.98, p = 0.005, Figures 4, S5). In general, males had lower FS (β = −0.44, p = 0.02) compared to females, and mice from 2016 had higher BCI (β = 1.09, p = 0.038) but lower FS (β = −0.63, p < 0.001) compared to 2015 (Figure S5). There were no effects of treatment detected on either metric of body condition, however among pregnant females, higher *H. polygyrus* infection intensity was associated with lower BCI scores (β = −0.64, p < 0.001) (Figure S5).

In the laboratory, supplemented diet resulted in higher total FS compared to control mice (β = 0.85, p = 0.019, but did not affect body mass (Figures S6–7). Males had both higher mass (β = 8.56, p < 0.001) and FS (β = 2.24, p < 0.001) compared to females (Figure S7). Lastly, higher *H. polygyrus* infection intensity was associated with overall decreased body mass (β = −1.40, p = 0.012, Figure S7).

### Supplemented nutrition increased total faecal IgA and *H. polygyrus-*specific IgG1

Total faecal IgA antibody concentrations differed between the years in the field experiment, with lower levels found in 2016 (β = −5.14, p = 0.0008). In 2016, wood mice on supplemented grids had significantly higher total faecal IgA antibody concentrations (β = 5.31 p = 0.007, no difference in 2015; Figure 5A; Table S8) and anthelminthic treatment was also associated with higher total faecal IgA in both years (β = 2.19, p = 0.031, Table S8). Body condition (BCI) was also found to be positively associated with higher concentrations of IgA (Figure 5A-B; Table S8) and was the only significant predictor of *H. polygyrus*-specific IgG1; where better body condition was associated with higher *H. polygyrus*-specific IgG1 concentration (β = 0.02, p = 0.01, Figure 5C; Table S8). Following controlled *H. polygyrus* infection in the laboratory, mice on a supplemented diet had significantly higher total faecal IgA 2-4 weeks post-infection (β = 2.40, p = 0.018) and *H. polygyrus-*specific IgG1 3 weeks post-infection (Figure 5D-E; Table S8).

**Figure 5.**
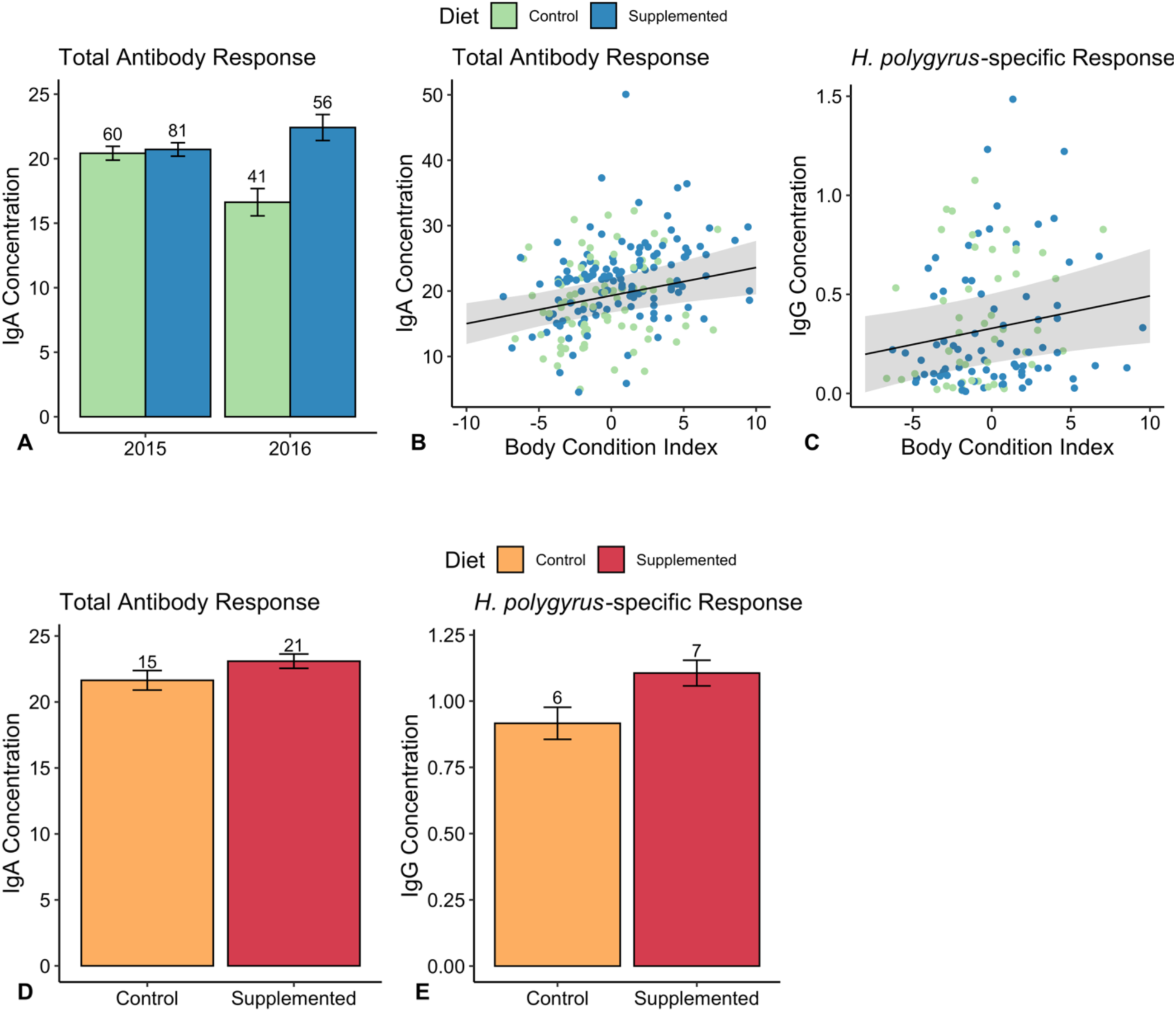
The impact of diet supplementation on wood mouse antibody responses, wild: top row; laboratory: bottom row. IgA Concentration (absolute, ng/uL) at all captures **(A)** across years and (B) compared to the body condition index (BCI). **(C)** *H. polygyrus* specific IgG1 concentration (standardised) at all captures compared to BCI. **(D)** Mean IgA Concentration (absolute, ng/uL) and (E) mean *H. polygyrus* specific IgG1 concentration (standardised) at days 14, 21, 28 post-primary challenge E. IgG1 Concentration d21 post-primary challenge. Bar plots represent raw means +/− SEM for the metric indicated. Numbers above bars indicate number of observations per group. Point-line plots represent raw data points and model-predicted regression slopes with 95% credibility interval ribbons.

## Discussion

We investigated the impact of a supplemented diet enriched with both micro- and macro-nutrients on helminth infection and immunity using a natural host-parasite relationship in both controlled laboratory and wild conditions. Our results demonstrate dramatic and surprisingly fast-acting benefits of supplemented nutrition for host resistance, immunity, treatment efficacy and body condition – effects which can be detected within just a few weeks in both settings. By conducting a parallel experiment with the same host and nematode species in controlled laboratory conditions we were able to overcome many of the limitations of field experiments; specifically, we controlled for variation among individuals in parasite exposure, demographic characteristics and nutritional status. We demonstrate that an enriched diet directly reduces both *H. polygyrus* infection and onward transmission in its natural host via improved host condition and immunity; suggesting that this type of integrated approach could be important for successful helminth management.

While the relationship between nutrition and gastrointestinal nematode infections have been extensively studied in the laboratory, the impact of this interaction is less well understood in natural populations. A previous field experiment on wood mice during winter found that supplementing diet with grass seeds led to a reduction in pinworms (*Syphacia stroma* and *Syphacia frederici*) but not cestodes (*Gallegoides arfaai* and *Hymenolepis diminuta*) or other nematodes (*Gongylonema neoplasticum*) [39]. Similarly, a recent study in laboratory mice released in semi-wild enclosures found no effect of restricted resource availability on *Trichuris muris* burden despite documenting increased host feeding behaviour, lower IL-13 expression, and lower *T. muris-*specific IgG1 production in these diet-restricted animals [52]. In addition, previous studies in wild systems that found benefits of supplementation on host condition and nematode infection in wild mouse populations (e.g. *Peromyscus spp.* [7] and *A. sylvaticus* [39]) mimicked food that was naturally available. In the present study, we found that experimentally supplementing naturally available resources with well-balanced food caused wood mice to rapidly and significantly reduce *H. polygyrus* infection burdens and transmission potential (as measured by egg shedding) compared to mice on control grids. The positive impacts of supplementation were observed after only 14 days, and were even detected in mice that visited the supplemented grids only briefly. This suggests resources are naturally limiting in wild wood mice populations, hampering their ability to produce a protective immune response to *H. polygyrus*.

Interestingly, we find very similar effects in the controlled laboratory experiment, with supplemented nutrition reducing worm burdens and egg shedding. This suggests that the benefits of whole-diet supplementation are not due to differences in wild wood mice behaviour or foraging patterns. Instead, we hypothesise they were driven by physiological and/or immunological changes. Further, the enriched diet reduced both *H. polygyrus* egg shedding during primary infection and susceptibility to reinfection compared to mice fed the standard, maintenance lab chow – the latter being similar to the diet used in many lab mouse studies. Protein and zinc levels have been previously found to impact susceptibility and protective immunity to *H. polygyrus* in lab mice [33,34,37]. However, much of this previous work was conducted using diet restrictions regimes (i.e. protein restriction (2-3% protein) compared to high protein (16-24% protein) [33,36,53–55]. We find similar and compelling effects of diet on infection in the laboratory but using more modest differences in macro-and micro-nutrients (i.e. 20% vs 14% protein, 80mg/kg vs 36mg/kg zinc) than in those studies. Our results suggest that diet-induced changes to nematode infection dynamics are not limited to severe malnutrition, nor to cases where there is a deficit in a specific nutrient. We hypothesise that the balance of multiple micro-and macro-nutrients contained in the enriched diet increased the host’s ability to generate a more protective immune response. However, more research is needed to determine if benefits of supplementation on condition and immunity to *H. polygyrus* are due to net effects of additional resources, or specific macro-and micro-nutrients contained in the supplemented diet.

Adequate levels of macro- and micro-nutrients are vital to normal function of cellular and humoral components of the immune system [21, 56] We found that our supplemented diet increased adaptive immunity in both wild and laboratory settings, specifically total faecal IgA and *H. polygyrus-*specific IgG1 levels were higher in supplemented individuals in the laboratory and total faecal IgA was higher in supplemented mice in 2016. Faecal IgA is an important component of resistance to gastrointestinal nematodes and has been used as an indicator of general gut health [57, 58], whereas parasite-specific IgG1 has a key role in the strong Th2 immune response induced by *H. polygyrus* [59]. These antibodies specifically play an important function in blocking the maturation of larvae into adult worms within the host intestinal tissue and reducing worm fecundity [60]. Our findings align with previous work suggesting that inadequate levels of nutrients (e.g. protein and zinc) compromise both general and specific host immune responses [33–35,52]. Although we saw weaker evidence for a direct effect of supplemented nutrition on antibody production in the wild, we found positive associations between the body condition index (BCI) and both total faecal IgA and the *H. polygyrus*-specific immune response. Therefore, the increase in body condition for supplemented individuals may indicate an indirect effect of supplementation on antibody levels and increased resistance. Typically, immune measures in the wild are difficult to interpret due to the context-dependency of immune phenotypes, and our limited ability to relate immune measures to exposure [47]. Thus, our results from our exposure-controlled colony experiment may be a more reliable indicator of how supplemented nutrition can impact both specific and general immune responses and impact helminth resistance.

Hosts in the wild are typically resource-limited, with finite energy to invest in immunity, reproduction, and other processes [22]. In addition to improving immunity, we found that a supplemented diet improves body condition and investment in reproduction. Mice, particularly pregnant females, on supplemented grids had increased host body condition scores (BCI; body condition index and FS; fat scores). Higher body condition during pregnancy, particularly for BCI which represents a body mass/ body length relative score, may indicate increased allocation to more or larger offspring. These results agree with previous research in which females in wood mouse populations supplemented with grass seeds during the winter bred earlier and had larger litter sizes [39]. In our laboratory experiment, supplementation had only a weak positive effect on fat scores, and did not affect BCI, contrary to previous laboratory studies which found that animals given lower protein had significantly lower weight, while our standard diet group did not lose weight over the experiment [33]. However, this may not be surprising, as wood mice in our laboratory colony are able to feed *ad libitum* and have higher body mass than those in the wild whom are likely to be chronically under some degree of dietary restriction.

Effective helminth control in endemic areas is difficult because even with readily available anthelmintic drugs, reinfection rates are usually high [11]. Limiting infection in a population requires (1) lowering worm burdens, (2) reducing onward transmission, and (3) preventing reinfection. In our wild population, we found that supplementation increased the efficacy of anthelmintic treatment, reducing *H. polygyrus* adult worm burdens and egg shedding to almost zero following treatment of mice on supplemented grids. This was consistent with supplemented individuals in the laboratory shedding no eggs during secondary challenge despite harbouring adult worms. Our results therefore indicate that nutrition can block transmission in addition to reducing parasite burdens. Further, we found that increased resistance to reinfection in supplemented individuals in the laboratory highlights an additional benefit of nutrition in managing nematodes in natural populations where post-treatment reinfection is commonplace. Given equivocal results from human trials investigating the addition of supplements with different nutrients [32] and the range of macro-and micro-nutrients implicated in immunity to gastrointestinal nematodes [55,61,62], our study presents key experimental results regarding the important role of balanced nutrition as a viable option for complementing helminth control interventions.

## Author Contributions

ARS, PAP, SAB and ABP designed the field studies. ARS, ABP and SAB conceived of the laboratory study. All authors were involved in the collection of field data. ARS, MEC, PAP, and SV performed the laboratory analyses. ARS performed the statistical analyses and wrote the first draft of the manuscript. All co-authors contributed to writing the final manuscript.

## Acknowledgements

This work was supported by PhD studentships from the Darwin Trust of Edinburgh to ARS and SV, a Torrance Bequest scholarship from the University of Edinburgh awarded to MC, a Wellcome Trust Institutional Strategic Support Fund (ISSF) grants to ABP (ISSF 2014; J22737) and SAB (097821/Z/11/Z), a targeted Institute of Biodiversity, Animal Health & Comparative Medicine Research Fellowship to SAB, a Wellcome Trust Strategic Grant for the Centre for Immunity Infection and Evolution (095831) to ABP, and a University of Edinburgh Chancellors Fellowship to ABP. We thank the Forestry Commission for permissions for field work in Callendar Park. *Heligmosomoides polygyrus* excretory-secretory product (HES) used in immunological assays was prepares and supplied by Rick Maizels (University of Glasgow) and Amy Buck (University of Edinburgh). Thanks also to Greg Albery for insightful comments on the manuscripts, and to the many tireless field assistants and volunteers involved in field and laboratory data collection. Lastly, we thank the EGLIDE (Edinburgh, Glasgow, and Liverpool Infectious Disease Ecology) group for their feedback and help with the analysis and interpretation of our results.

## Supplementary Material: Sweeny *et al*. 2019 Supplemented nutrition increases immunity against helminths

## Methods

Here we provide additional information on methods used in the field and in the laboratory experiment, as well as rationale for approach to age and body condition analysis [1].

### Field Experiment

#### Experimental trapping regime

In 2015, we live-trapped three grids (1 supplemented grid, 2 control grids, 49 trapping stations per grid with 2 traps/station, 10m between each trap, for a total area of 3600m^2^), while in 2016, we trapped four grids (2 supplementation and 2 control grids; with each grid set up as a 6×5 array of 30 trapping stations with 2 traps/station, 10m between each trap, for a total area of 2000m^2^). All grids in both years were spaced a minimum of 50m from each other to minimise mouse movement between grids, and grids were randomly assigned to nutrition regimes prior to the start of the experiment. During trapping, each trap was set with cotton wool bedding, and was baited with seeds, carrot, mealworms, and TransBreed™ pellets (on supplemented grids only), set in the early evening (16.00-18.00) and then checked early the following morning.

#### Host data and sampling

At each capture, weight in grams and length in mm was measured for each individual. Body fat scores were assessed on a scale of 1-5 (emaciated-obese) by palpating the sacroiliac bones (back and pubic bones) as detailed in [2]. After field data collection, body condition index (BCI) was calculated by obtaining the residuals of an ordinary least squares (OLS) regression of mass against length [3]. We did not calculate body condition index for the colony mice as those measures would be obscured by the fact that they are, on average, much heavier compared to their wild counterparts. Therefore, condition in the laboratory was measured using mass and total fat score only.

Sex and reproductive status were assigned by visual examination of the genitals as male wood mice have a greater urogenital distance than females. Males were classed into the following reproductive categories: abdominal (testes non-visible); descended, or scrotal. Females were classed into the following reproductive categories: non-perforate vagina; perforate vagina; pregnant or lactating. Animals which are scrotal, pregnant, or lactating are considered reproductive for binary reproductive status assignment. Age of mice (classed as: juvenile, subadult, adult) was determined by weight and coat colour based on juvenile moulting patterns. Generally, juveniles weigh 10g or less, subadults weigh between 10-15g, and adults are 15g or heavier. Juveniles have a distinctly different coat colour (grey, compared to brown colour of adults), while subadults have an intermediate colour coat (grey/brown). Where available (for sacrificed animals), eyes were collected and stored in 10% formalin for dissection of eye lenses for assessment of animal age. Eye lens mass has been shown to strongly correlate with age in many species (rodents and others), and has successfully been used to distinguish age classes for both laboratory and wild mice [4,5]. Eyes collected from sacrificed animals were removed from their container and left at room temperature for 5-10 minutes to allow the formalin to evaporate. Eye lenses were then extracted and dried at 70°C overnight. They were then weighed to the nearest mg using a precision balance. The combined weight of both eye lenses (log-transformed) for each individual were used as a proxy for age. We calculated the relationship between age and eye lens weight using wood mice of known ages from our colony to be (Figure S2):

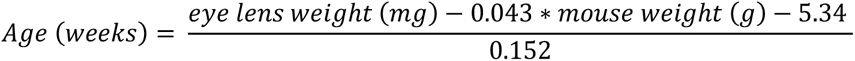

Blood samples collected in the field were centrifuged at 12,000 rpm for 10 minutes for collection of the sera and then stored at −80°C upon returning to the laboratory. Faecal samples were collected from previously sterilised traps and then preserved in 10% formalin at 4°C until processing and 2-3 pellets from each faecal sample were stored at −80°C for faecal IgA antibody measures. Small intestine, caecum, and colon of each sacrificed individual were stored in 1X phosphate-buffered saline and dissected on the same day for counts of adult *H. polygyrus* worms present.

### Laboratory experiment

#### Wood mouse colony

We maintain a formerly-wild, but now lab-reared wood mouse colony in standard laboratory conditions at the University of Edinburgh. The colony has been in captivity for many generations, but the wood mice are purposely outbred to maintain genetic diversity. In this experiment, all mice were housed individually in ventilated cages (Techniplast, 1285L) with food and water ad libitum. *H. polygyrus* L3 larvae were isolated from the same Callendar Wood wild wood mouse population and were screened using PCR diagnostics to ensure the isolate was not contaminated with any other known mouse parasites or pathogens (IDEXX Bioresearch, Germany), and then passaged several times through colony-housed wood mice [6].

#### Diet Information

Wood mice in the colony were fed two different formulations of standard laboratory rodent chow. Control mice were fed with standard maintenance chow (RM1™), whereas supplemented mice were fed a specially-formulated chow (TransBreed™), which has been designed to include higher fat, protein, and micronutrient contents (Table SI1). We selected those diets to approximate control and supplemented nutrition in our field experiment as closely as possible. Both diets were fed ad libitum and provided adequate maintenance nutrition

**Table S1.**
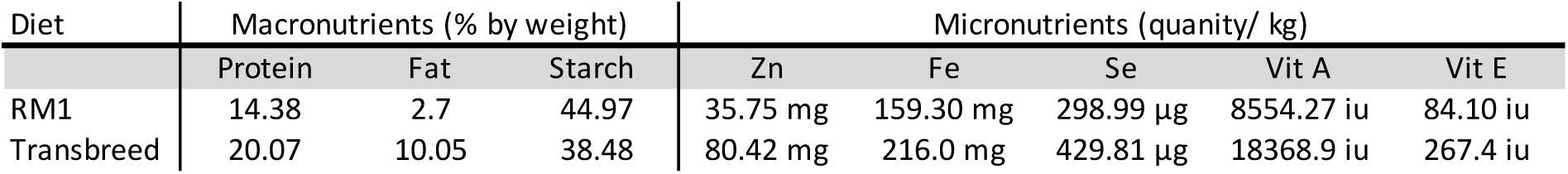
Nutrition content of SDS chow used in laboratory experiment comparing *H. polygyrus* infection for mice on both a standard (RM1™) and supplemented (Transbreed™) diet. Micronutrients included are selected elements and vitamins which have previously been implicated in host response to helminth infection, however full calculated analysis of RM1™ and Transbreed™ nutritional content can be found at http://www.sdsdiets.com/pdfs/RM1P-E-FG.pdf & http://www.sdsdiets.com/pdfs/TransBreed.pdf.

#### Parasite material & anthelmintic treatment

Third stage larvae (L3) of *H. polygyrus* were originally derived from wild wood mice from Callendar Park that were infected with *H. polygyrus* [6]. Since then they have been passaged approximately ten times through colony-housed wood mice at the University of Edinburgh. In order to extract *H. polygyrus* eggs from faecal samples, the pellets were broken up and mixed with inactivated charcoal to mimic soil. The charcoal-soil mix was spread thinly on moist filter paper maintained in petri dishes at 17C. Larvae started to hatch after approximately 12 days and were collected into PBS/water? (cannot remember anymore) and kept at 4C until use. Prior to infections, larvae concentrations were adjusted to a final concentration of 200 L3/ 150uL. Infective doses were administered to mice via oral gavage.

Ivermectin and Pyrantel pamoate are broad-spectrum anthelmintics which target adult and larval stages (Ivermerctin) and adult stages (Pyrantel) of *H. polygyrus* in both laboratory and wild mice [7-9]. Previous work in wild *A. sylvaticus* found that the combination of Ivermectin and Pyrantel at 9.4mg/kg and 100 mg/kg, respectively, efficiently cleared *H. polygyrus* infection for ∼14 days [10].

#### Sampling

Faecal samples in the colony were collected by changing the cage bedding ∼12 hr prior to each collection and then collecting faecal pellets from the freshly-used bedding and then preserving the samples in 10% buffered formalin. A small sample of 2-3 pellets was also collected to measure faecal IgA.

### Laboratory assays

#### Parasite counts

Saturated salt solution was added to formalin-preserved faecal samples to concentrate eggs on a coverslip, counted at 10X magnification, and adjusted by sample weight to the number of eggs per gram faeces (EPG). Adult *H. polygyrus* were counted from PBS-preserved small intestine sections within 5 hours of dissection.

#### Antibody assays

ELISAs were performed to measure (1) total faecal IgA concentration and (2) serum *H. polygyrus-* specific IgG1 titres for each mouse at each capture/sampling point. For IgA ELISA, a 3:1 volume of protease inhibitor solution (Complete Mini Protease Inhibitor Tablets, Roche) was added each faecal sample and then homogenised. These faecal extractions were then incubated for 1hr at room temperature and then centrifuged at 12,000 rpm for 5min. Supernatants were separated from faecal pellets and stored at −80C. 96-well microplates (Nunc™ MicroWell™, ThermoScientifc™) were coated with goat anti-mouse IgA (Southern Biotech 1040-01, 2ug/ml) diluted in carbonate buffer overnight at 4C. Capture antibody was then flicked off plates and 4% BSA-TBS was added and incubated for 2hr at 37C to block non-specific binding sites. Faecal extracts were diluted 1:50 in 1% BSA TBS in triplicate and added to plates and incubated overnight at 4C. Two serial dilutions of IgA standard (BD Bioscience 3039828) were included on each plate as positive controls and to obtain curves for calculation of IgA concentration. Following incubation, plates were washed 3 times with TBS-Tween and 50uL of goat anti-mouse IgA-HRP (Southern Biotech 1040-05) diluted 1:4000 in 1% BSA TBS was added to each well. Plates were incubated for 1hr and washed 4x with TBS-Tween and 2x with ddH20. 50ul TMB substrate was added to each well and plates were developed for 7 minutes protected from light. After that, 50uL of 0.18M sulphuric acid was added to each well to stop the enzymatic reaction. Plates were read at 450nm and sample concentrations were determined by fitting a 4-parameter logistic regression to standard curves.

To determine *H. polygyrus-*specific IgG1 titres from blood sera, 96-well microplates were coated with *H. polygyrus* excretory-secretory antigen (HES, 1.0ug/ml; obtained from Amy Buck, University of Edinburgh, and Rick Maizels, University of Glasgow) diluted in carbonate buffer overnight at 4C. Capture antigen was then flicked off plates and 4% BSA-TBS was added and incubated for 2hr at 37C to block non-specific binding sites. Sera samples were prepared as twofold serial dilutions with a starting concentration of 1:100in 1% BSA TBS and added to plates, and incubated overnight at 4C. Two serial dilutions consisting of sera from *Mus musculus domesticus* experimentally infected with *H. polygyrus* in the laboratory were included on each plate as positive controls. Following incubation, plates were washed 3x with TBS-Tween and 50uL of goat anti-mouse IgG1-HRP (Southern Biotech 1070-05) diluted 1:2000 in 1% BSA TBS was added to each well. Plates were incubated for 1hr and washed 4x with TBS-Tween and 2x with ddH20. 50ul TMB substrate was added to each well and plates were developed for 7min protected from light, at which point 50uL of 0.18M sulphuric acid was added to each well to stop the enzymatic reaction. Plates were prepared with serial dilutions of reference and experimental samples, and a dilution factor of 1:200 was selected for calculation of relative antibody concentrations. Standardised IgG1 concentrations were calculated by plate as follows:

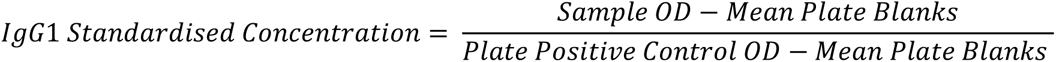

We assigned a value of 0 to samples for which the OD did not exceed 3x SD of control blanks as we considered them indistinguishable from no IgG1 response. We refer to both IgA and IgG1 values as ‘antibody concentration’.

**Table S2.** Model formulae for analyses of wild and laboratory experiments. EPG stands for eggs per gram faeces; dpi for days post-infection.

| MODEL GROUP | SYSTEM | RESPONSE | EXPERIMENT TIMEPOINT | MODEL CLASS | MODEL FAMILY | FIXED EFFECTS* | INTERACTIONS | RANDOM EFFECTS |
| --- | --- | --- | --- | --- | --- | --- | --- | --- |
| **H. POLYGYRUS INFECTION** | Wild | EPG | First capture | GLM | Negative Binomial | Body Mass + Reproductive status + Sex + Diet |  | Grid*Year |
|  |  | EPG (mean) | Trapping duration | GLM |  | as above + Treatment | Diet:Treatment | Grid*Year |
|  |  | Adult worm | End point | GLM |  | as above + Age | Diet:Treatment | Grid*Year |
|  | Lab | EPG (peak) | Primary challenge | GLM | Negative Binomial | Body Mass+ Sex + Age + Group+ Diet |  |  |
|  |  | EPG (total) | Primary challenge |  |  | as above |  |  |
|  |  | Adult worm | End point |  |  | as above | Diet:timepoint |  |
| **ANTIBODY RESPONSE** | Wild | IgA (total) | All captures | GLMM | Gaussian | Body condition index + Reproductive status+ Sex, H. polygyrus infection + Diet + Year | Diet:Year | ID + Grid*Year |
|  |  | IgG1 (specific) |  |  |  | as above |  | Grid*Year |
|  | Lab | IgA (total) | 14 & 21 dpi | GLMM | Gaussian | Sex + Age + H. polygyrus infection + Experiment Timepoint + Diet | Diet:timepoint | ID |
|  |  | IgG1 (specific) | 21 dpi | GLM |  | Sex + Age + H. polygyrus infection (n worms) + Diet |  |  |
| **BODY CONDITION** | Wild (a) adult, non-pregnant | BCI | All captures | LMM | Gaussian | Reproductive status + Sex + H. polygyrus infection (Log, EPG) + Year + Day + Diet + Treatment | Diet:Day | ID + Grid*Year |
|  |  | Total fat score |  |  |  | as above | Diet:Day | ID + Grid*Year |
|  | Wild (a) pregnant | BCI |  |  |  | H. polygyrus infection (Log, EPG) + Year + Day + Diet + Treatment | Diet:Day | ID + Grid*Year |
|  |  | Total fat score |  |  |  | as above | Diet:Day | ID + Grid*Year |
|  | Lab | Body mass (g) | All captures | LMM | Gaussian | Sex + Age + H. polygyrus infection (Log, EPG) + Day + Diet | Diet:Day | ID |
|  |  | Total fat score |  |  |  | as above | Diet:Day | ID |

**Table S3.**
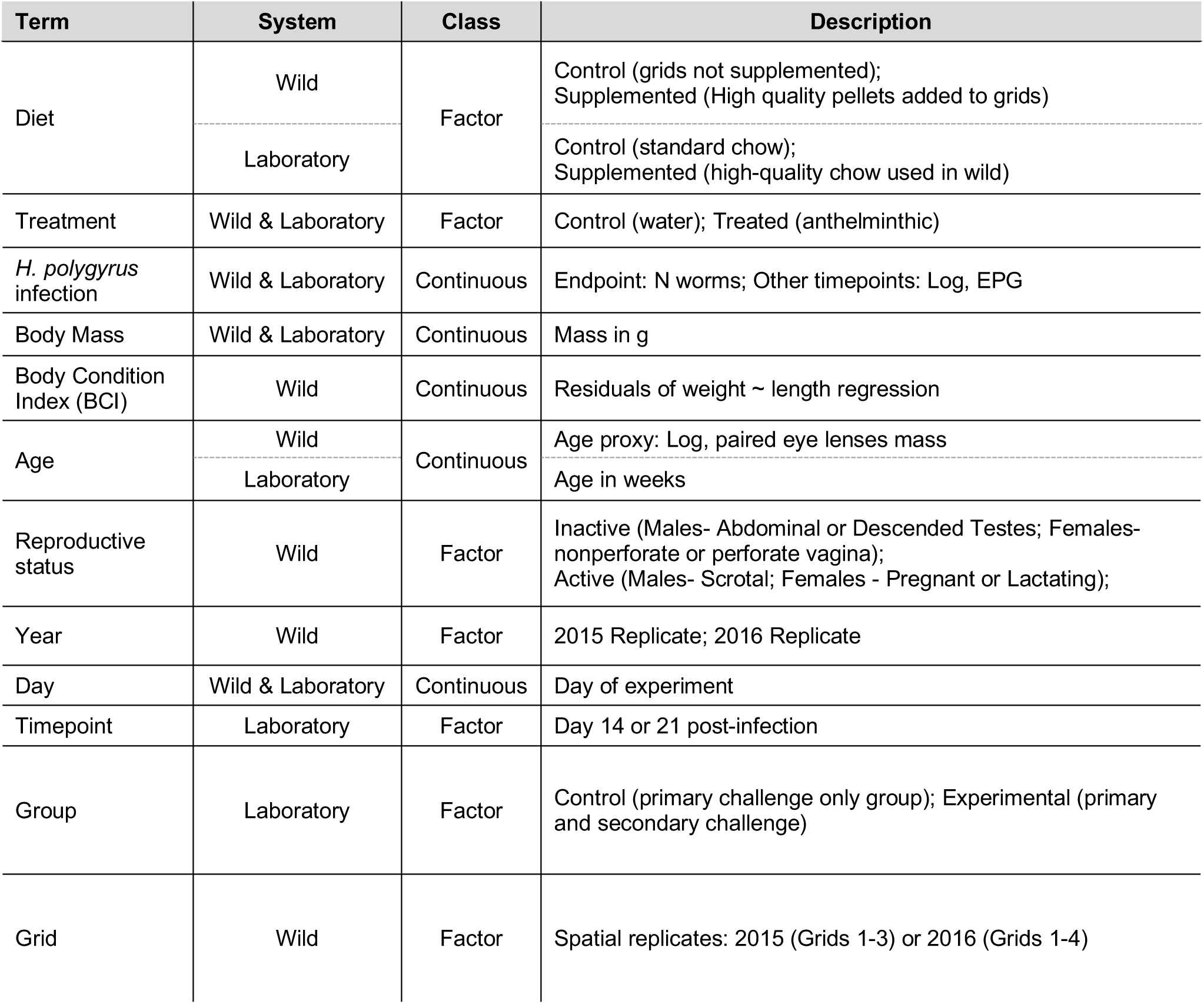
Description of fixed effects included in models

| Term | System | Class | Description |
| --- | --- | --- | --- |
| Diet | Wild | Factor | Control (grids not supplemented);<br>Supplemented (High quality pellets added to grids) |
|  | Laboratory |  | Control (standard chow);<br>Supplemented (high-quality chow used in wild) |
| Treatment | Wild & Laboratory | Factor | Control (water); Treated (anthelminthic) |
| *H. polygyrus* infection | Wild & Laboratory | Continuous | Endpoint: N worms; Other timepoints: Log, EPG |
| Body Mass | Wild & Laboratory | Continuous | Mass in g |
| Body Condition Index (BCI) | Wild | Continuous | Residuals of weight ~ length regression |
| Age | Wild | Continuous | Age proxy: Log, paired eye lenses mass |
|  | Laboratory |  | Age in weeks |
| Reproductive status | Wild | Factor | Inactive (Males- Abdominal or Descended Testes; Females- nonperforate or perforate vagina);<br>Active (Males- Scrotal; Females - Pregnant or Lactating); |
| Year | Wild | Factor | 2015 Replicate; 2016 Replicate |
| Day | Wild & Laboratory | Continuous | Day of experiment |
| Timepoint | Laboratory | Factor | Day 14 or 21 post-infection |
| Group | Laboratory | Factor | Control (primary challenge only group); Experimental (primary and secondary challenge) |
| Grid | Wild | Factor | Spatial replicates: 2015 (Grids 1-3) or 2016 (Grids 1-4) |

## Raw Data Summary, Full Model Output & Model Variation Output

This section includes raw data summaries and full model output for further interpretation of data presented in the main text. Additionally, we describe the models using a 3-level factor for supplementation in the wild to more accurately classify mice who were captured on both supplemented and control grids. Model output for full main models is shown in Figure 3. For model variation described, fixed effects are held the same as in the main models, with the exception of the addition of Supplement category (‘Mix’) (Figure S1). Estimates for fixed effects are compared between main models and their variations.

For both models where an effect of supplemented nutrition was detected, both levels of supplemented nutrition (mix and supplemented) lowered *H. polygyrus* EPG and worm burden, and the magnitude and direction of other fixed effects closely mirrored those of the base model of comparison (Figures S1& S4).

**Table S4.** Raw data *H. polygyrus* summary. Data represents mean eggs/gram (EPG) or number of worms (± SE), and sample sizes for the indicated group.

|  |  | Control (dH<sub>2</sub>O) |  | Anthelmintic |  |
| --- | --- | --- | --- | --- | --- |
|  |  | Control | Supplemented | Control | Supplemented |
| *Wild* | First Capture (mean EPG) | 23.55 (± 8.72);<br>n=36 | 12.41 (± 3.20);<br>n=52 | na | na |
| | Trapping During (mean EPG) | 43.68 (± 8.78);<br>n=37 | 29.96 (± 11.07);<br>n=46 | 29.02 (± 19.99);<br>n=36 | 0.136 (± 0.136);<br>n=48 |
| | End point (Number of adult worms) | 30.50 (± 7.99);<br>n=6 | 12.21 (± 3.96);<br>n=14 | 1.71 (± 0.89);<br>n=7 | 0.11 (± 0.11);<br>n=9 |

|  | Group | Primary Challenge |  | Secondary Challenge |  |
| --- | --- | --- | --- | --- | --- |
|  |  | Control | Supplemented | Control | Supplemented |
| *Colony* | Peak EPG | 48.03 (± 25.79);<br>n=9 | 19.59 (± 8.42);<br>n=10 | 1.76 (± 1.19);<br>n=6 | 0.00 (± 0.00);<br>n=7 |
| | Sum EPG | 63.86 (± 26.82);<br>n=9 | 28.26 (± 11.38);<br>n=10 | 2.72 (± 2.07);<br>n=6 | 0.00 (± 0.00);<br>n=7 |
| | End point (Number of adult worms) | 9.67 (± 3.38);<br>n=3 | 12.33 (± 4.91);<br>n=3 | 28.33 (± 6.50);<br>n=6 | 6.71 (± 1.98);<br>n=7 |

**Table S5.** Model estimates for fixed effects on wild *H. polygyrus* infection.

| Term | First Capture<br>EPG |  |  | Trapping Duration<br>EPG |  |  | End Point<br>Number of adult worms |  |  |
| --- | --- | --- | --- | --- | --- | --- | --- | --- | --- |
|  | Estimate (CI) | Std.error | p.value | Estimate (CI) | Std.error | p.value | Estimate (CI) | Std.error | p.value |
| Diet, Supplemented:<br>Treated, Anthelmintic |  |  |  | -4.51 (-8.12 - -0.9) | 1.84 | **0.014** | -2.25 (-5.26 - 0.75) | 1.53 | 0.142 |
| Treated, Anthelmintic |  |  |  | -2.11 (-4.58 - 0.35) | 1.26 | 0.092 | -2.74 (-4.26 - -1.22) | 0.78 | **< 0.001** |
| Diet, Supplemented | -1.42 (-2.59 - -0.24) | 0.6 | **0.018** | -1.56 (-3.65 - 0.53) | 1.07 | 0.145 | -1.2 (-2.38 - -0.03) | 0.6 | **0.045** |
| Sex, Male | -0.99 (-2.27 - 0.3) | 0.66 | 0.132 | 0.25 (-1.59 - 2.09) | 0.94 | 0.787 | -1.16 (-2.58 - 0.26) | 0.72 | 0.109 |
| Reproductive, active | -0.53 (-1.74 - 0.68) | 0.62 | 0.387 | 1.37 (-0.26 - 3.01) | 0.83 | 0.100 | -0.25 (-1.26 - 0.77) | 0.52 | 0.632 |
| Body Mass, g | 0.15 (-0.01 - 0.32) | 0.08 | 0.065 | 0.3 (0 - 0.6) | 0.15 | 0.053 | 0.21 (0.03 - 0.39) | 0.09 | **0.019** |
| Age<br>(Eye lens weight, mg) |  |  |  |  |  |  | 2.22 (0.41 - 4.04) | 0.93 | **0.016** |
| (Intercept) | 1.42 (-1.32 - 4.16) | 1.4 | 0.309 | -2.46 (-7.84 - 2.91) | 2.74 | 0.369 | -6.32 (-11.48 - -1.16) | 2.63 | **0.016** |

**Table S6.** Model estimates for fixed effects on laboratory *H. polygyrus* infection

|  | Primary Challenge<br>Peak EPG |  |  | Primary Challenge<br>Sum EPG |  |  | End Point<br>(Primary & Secondary Challenge)<br>Number of adult worms |  |  |
| --- | --- | --- | --- | --- | --- | --- | --- | --- | --- |
| Term | Estimate (CI) | Std.error | p.value | Estimate (CI) | Std.error | p.value | Estimate (CI) | Std.error | p.value |
| Diet, Supplemented:<br>Secondary Challenge |  |  |  |  |  |  | -1.76 (-2.88 - -0.64) | 0.57 | **0.002** |
| Diet, Supplemented | -1.09 (-1.99 - -0.19) | 0.46 | **0.017** | -1.07 (-2.04 - -0.11) | 0.49 | **0.03** | 0.21 (-0.73 - 1.14) | 0.48 | 0.663 |
| Group, Experimental | 1.96 (0.86 - 3.05) | 0.56 | **<0.001** | 2.22 (1.07 - 3.37) | 0.59 | **<0.001** | 0.89 (0.07 - 1.71) | 0.42 | **0.033** |
| Sex, Male | -0.51 (-2.05 - 1.03) | 0.79 | 0.515 | 0.06 (-1.62 - 1.74) | 0.86 | 0.945 | -0.08 (-1 - 0.83) | 0.47 | 0.857 |
| Body Mass, g | 0.01 (-0.17 - 0.19) | 0.09 | 0.902 | -0.06 (-0.25 - 0.14) | 0.1 | 0.57 | 0.06 (-0.04 - 0.16) | 0.05 | 0.248 |
| Age (Weeks) | -0.01 (-0.09 - 0.08) | 0.04 | 0.875 | -0.01 (-0.1 - 0.08) | 0.05 | 0.81 | -0.02 (-0.08 - 0.04) | 0.03 | 0.428 |
| (Intercept) | 2.82 (-0.9 - 6.54) | 1.9 | 0.138 | 3.52 (-0.46 - 7.5) | 2.03 | 0.083 | 1.84 (-0.53 - 4.21) | 1.21 | 0.127 |

**Table S7.** Model estimates for fixed effects on body condition in wild wood mice

| Term | All Adults |  |  |  |  |  | Pregnant Females |  |  |  |  |  |
| --- | --- | --- | --- | --- | --- | --- | --- | --- | --- | --- | --- | --- |
|  | Body Condition Index |  |  | Total Fat Score |  |  | Body Condition Index |  |  | Total Fat Score |  |  |
|  | Estimate (CI) | Std. error | p.value | Estimate (CI) | Std. error | p.value | Estimate (CI) | Std. error | p.value | Estimate (CI) | Std. error | p.value |
| Diet, Supplemented: Day | -0.16<br>(-0.28 - -0.04) | 0.06 | **0.011** | -0.09<br>(-0.15 - -0.02) | 0.03 | **0.007** | -0.61<br>(-0.76 - -0.45) | 0.08 | **<0.001** | -0.15<br>(-0.3 - 0) | 0.08 | **0.045** |
| Diet, Supplemented | 1.46<br>(0.34 - 2.59) | 0.57 | **0.011** | 0.64<br>(0.18 - 1.11) | 0.24 | **0.006** | 8.32<br>(5.5 - 11.15) | 1.44 | **<0.001** | 1.98<br>(0.6 - 3.37) | 0.71 | **0.005** |
| Day | -0.03<br>(-0.13 - 0.07) | 0.05 | 0.525 | 0.01<br>(-0.04 - 0.07) | 0.03 | 0.582 | 0<br>(-0.11 - 0.12) | 0.06 | 0.969 | 0.11<br>(-0.01 - 0.22) | 0.06 | **0.062** |
| Year, 2016 | 1.02<br>(0.04 - 1.99) | 0.5 | **0.041** | -0.81<br>(-1.14 - -0.47) | 0.17 | **<0.001** | 1.57<br>(-0.8 - 3.95) | 1.21 | 0.194 | -0.83<br>(-1.72 - 0.06) | 0.46 | **0.069** |
| Log, *H. polygyrus* EPG | 0.08<br>(-0.14 - 0.29) | 0.11 | 0.49 | 0.03<br>(-0.07 - 0.12) | 0.05 | 0.583 | -0.64<br>(-0.87 - -0.41) | 0.12 | **<0.001** | 0.06<br>(-0.2 - 0.32) | 0.13 | 0.64 |
| Treated, Anthelmintic | 0.28<br>(-0.71 - 1.26) | 0.5 | 0.582 | -0.14<br>(-0.49 - 0.21) | 0.18 | 0.429 | 0.26<br>(-2.27 - 2.79) | 1.29 | 0.838 | 0.16<br>(-0.81 - 1.13) | 0.5 | 0.746 |
| Sex, Male | 0.57<br>(-0.42 - 1.55) | 0.5 | 0.258 | -0.25<br>(-0.59 - 0.09) | 0.17 | **0.15** |  |  |  |  |  |  |
| Reproductive, active | 1.53<br>(0.74 - 2.33) | 0.4 | **<0.001** | -0.22<br>(-0.57 - 0.13) | 0.18 | **0.215** |  |  |  |  |  |  |
| Intercept | -2.6<br>(-4.03 - -1.17) | 0.73 | **<0.001** | 5.98<br>(5.41 - 6.56) | 0.29 | **<0.001** | -0.46<br>(-4.24 - 3.32) | 1.93 | 0.81 | 4.57<br>(2.81 - 6.34) | 0.9 | **<0.001** |

**Table S8.**
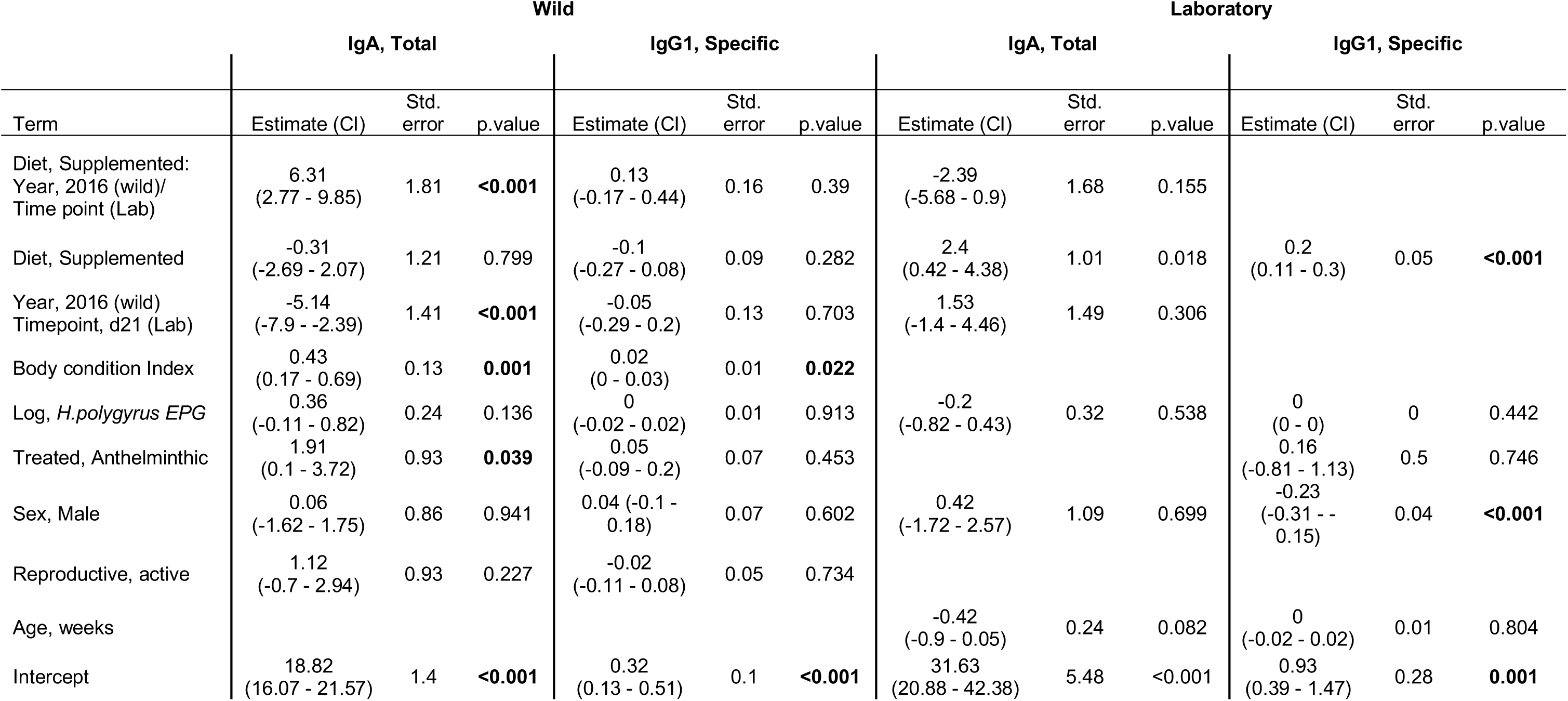
Model estimates for fixed effects on immune responses in wild and laboratory wood mice.

| Term | Wild |  |  |  |  |  | Laboratory |  |  |  |  |  |
| --- | --- | --- | --- | --- | --- | --- | --- | --- | --- | --- | --- | --- |
|  | IgA, Total |  |  | IgG1, Specific |  |  | IgA, Total |  |  | IgG1, Specific |  |  |
|  | Estimate (CI) | Std. error | p.value | Estimate (CI) | Std. error | p.value | Estimate (CI) | Std. error | p.value | Estimate (CI) | Std. error | p.value |
| Diet, Supplemented:<br>Year, 2016 (wild)/<br>Time point (Lab) | 6.31<br>(2.77 - 9.85) | 1.81 | **<0.001** | 0.13<br>(-0.17 - 0.44) | 0.16 | 0.39 | -2.39<br>(-5.68 - 0.9) | 1.68 | 0.155 |  |  |  |
| Diet, Supplemented | -0.31<br>(-2.69 - 2.07) | 1.21 | 0.799 | -0.1<br>(-0.27 - 0.08) | 0.09 | 0.282 | 2.4<br>(0.42 - 4.38) | 1.01 | 0.018 | 0.2<br>(0.11 - 0.3) | 0.05 | **<0.001** |
| Year, 2016 (wild)<br>Timepoint, d21 (Lab) | -5.14<br>(-7.9 - -2.39) | 1.41 | **<0.001** | -0.05<br>(-0.29 - 0.2) | 0.13 | 0.703 | 1.53<br>(-1.4 - 4.46) | 1.49 | 0.306 |  |  |  |
| Body condition Index | 0.43<br>(0.17 - 0.69) | 0.13 | **0.001** | 0.02<br>(0 - 0.03) | 0.01 | **0.022** |  |  |  |  |  |  |
| Log, *H. polygyrus* EPG | 0.36<br>(-0.11 - 0.82) | 0.24 | 0.136 | 0<br>(-0.02 - 0.02) | 0.01 | 0.913 | -0.2<br>(-0.82 - 0.43) | 0.32 | 0.538 | 0<br>(0 - 0) | 0 | 0.442 |
| Treated, Anthelmintic | 1.91<br>(0.1 - 3.72) | 0.93 | **0.039** | 0.05<br>(-0.09 - 0.2) | 0.07 | 0.453 |  |  |  | 0.16<br>(-0.81 - 1.13) | 0.5 | 0.746 |
| Sex, Male | 0.06<br>(-1.62 - 1.75) | 0.86 | 0.941 | 0.04 (-0.1 - 0.18) | 0.07 | 0.602 | 0.42<br>(-1.72 - 2.57) | 1.09 | 0.699 | -0.23<br>(-0.31 - -0.15) | 0.04 | **<0.001** |
| Reproductive, active | 1.12<br>(-0.7 - 2.94) | 0.93 | 0.227 | -0.02<br>(-0.11 - 0.08) | 0.05 | 0.734 |  |  |  |  |  |  |
| Age, weeks |  |  |  |  |  |  | -0.42<br>(-0.9 - 0.05) | 0.24 | 0.082 | 0<br>(-0.02 - 0.02) | 0.01 | 0.804 |
| Intercept | 18.82<br>(16.07 - 21.57) | 1.4 | **<0.001** | 0.32<br>(0.13 - 0.51) | 0.1 | **<0.001** | 31.63<br>(20.88 - 42.38) | 5.48 | **<0.001** | 0.93<br>(0.39 - 1.47) | 0.28 | **0.001** |

**Figure S2.**
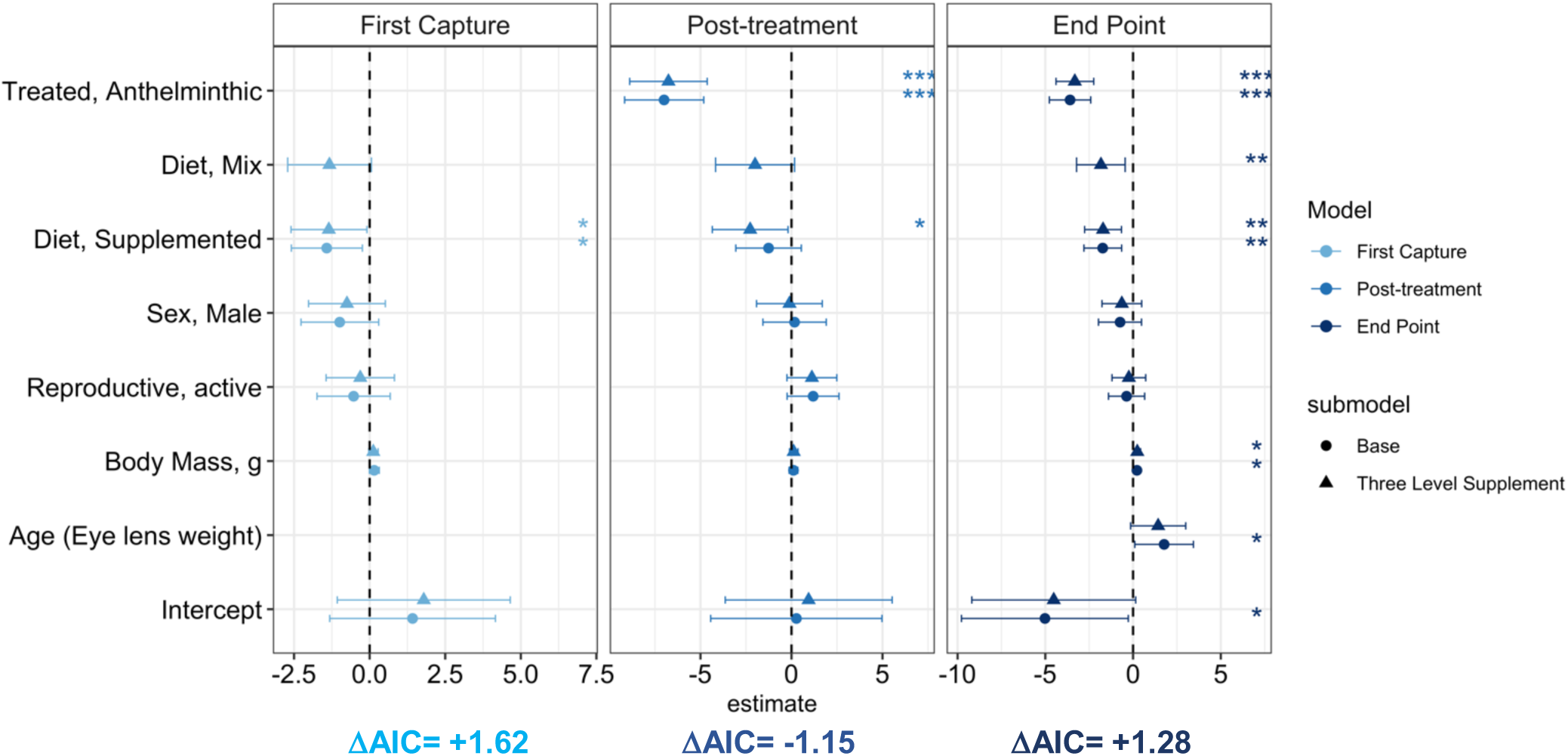
Effect size estimates from models investigating the effect of supplemented nutrition on *H. polygyrus* infection, accounting for individuals captured only a portion of time on supplemented grids (mix). Panels represent separate models for first capture, infection abundance (EPG); average EPG across two weeks for individuals captured beyond first capture and after assignment to treatment categories; end point burden (adult worm count) for individuals culled 12-16 days post first capture. Models represented are identical to those included in Fig 3 with the exception of the inclusion of an additional factor level in the supplemented nutrition explanatory variable. Points and ranges represent model estimates and 95% credibility estimates for each model. Asterisks indicate the significance of variables: ***, ** and * indicate P<0.001, P<0.01 and P<0.05 respectively. Change in AIC from main models is included in the bottom left of each panel.

## Additional figures

**Figure S3.**
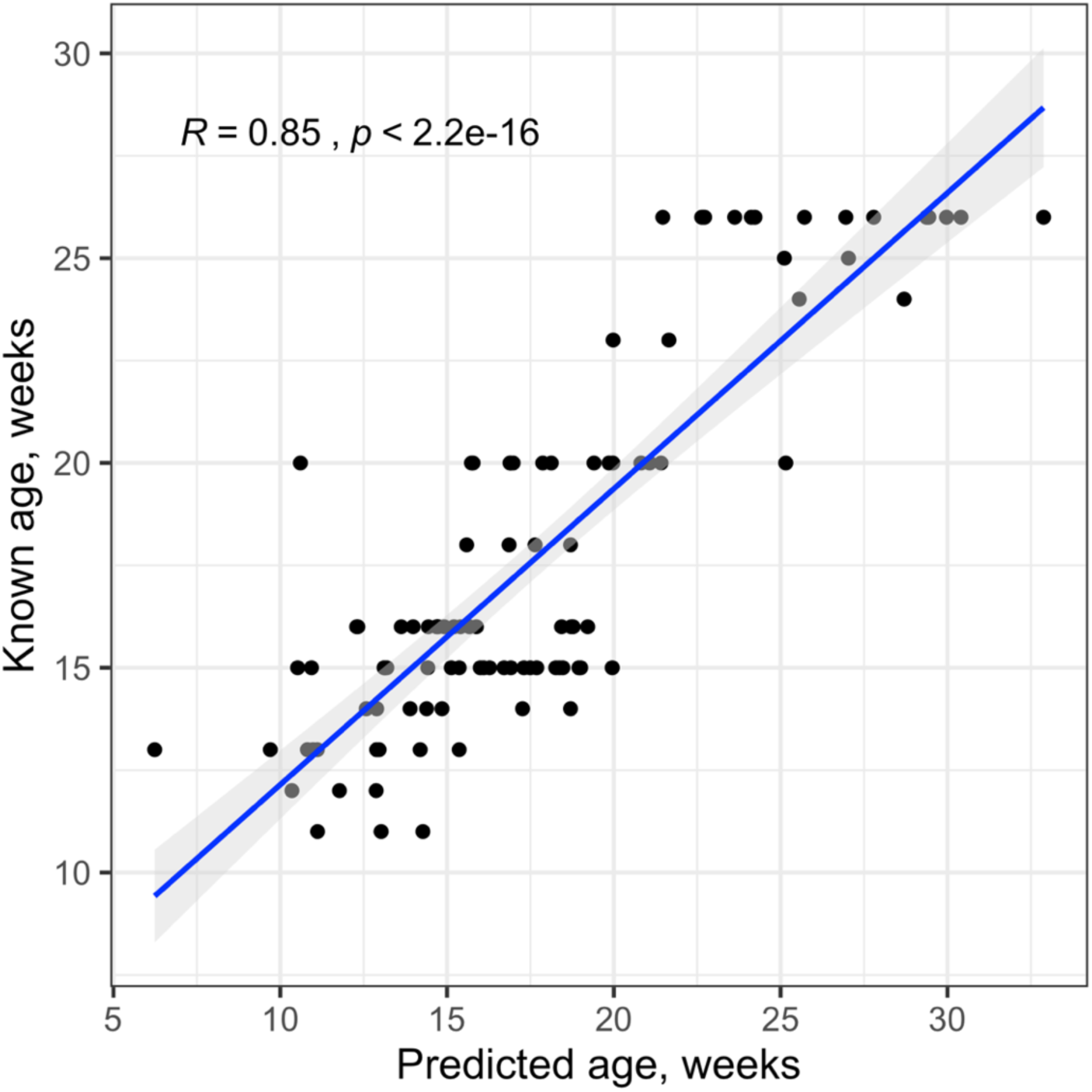
Correlation of wood mouse age predicted from eye lens weight and known age (in weeks) from colony wood mice. Pearson’s R with 95% credibility intervals and significance of the correlation is included for both 2015 and 2016 data. Points have been jittered by 10% of raw values to aid in visualisation.

**Figure S4.**
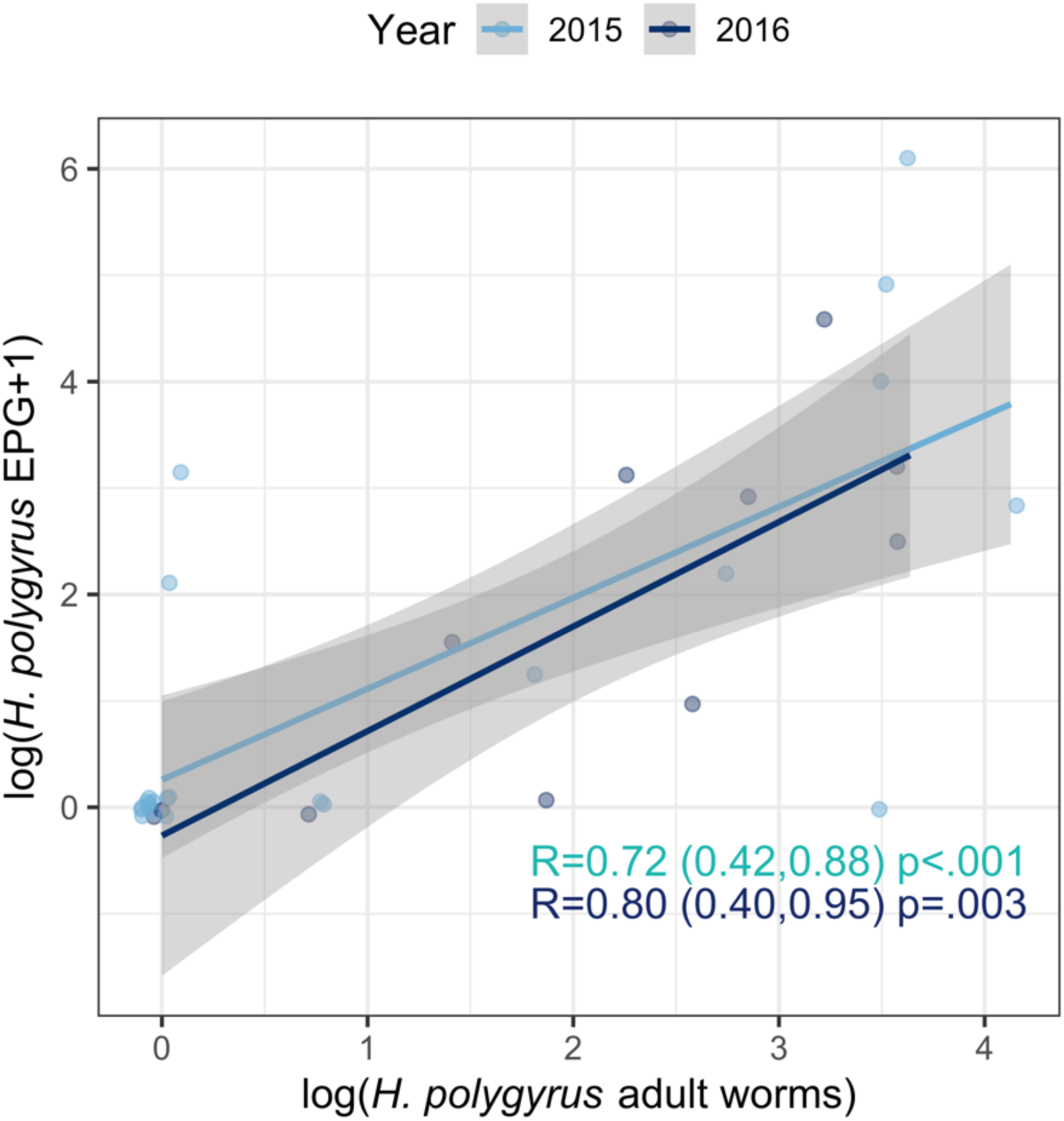
Correlation between *H. polygyrus* eggs/ gram faeces (EPG) and adult worm burden from culled animals where data was available for both metrics (N_i_=36). Both values are log-transformed (EPG or count +1). Pearson’s R with 95% credibility intervals and significance of the correlation is included for both 2015 and 2016 data. Points have been jittered by 10% of raw values to aid in visualisation.

**Figure S5.**
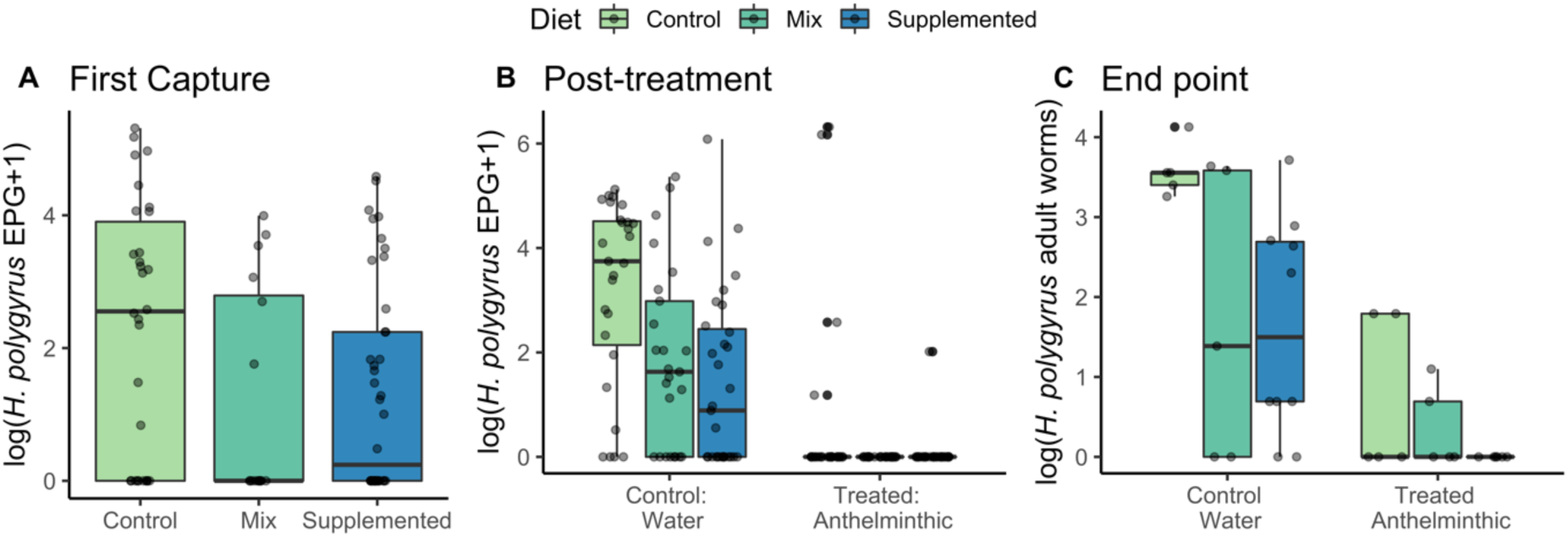
Effect of supplemented nutrition on H. polygyrus infection in wild wood mice, accounting for both individuals who were found exclusively on supplemented grids or individuals who were found on supplemented grids only a portion of the time (‘Mix”). A. Infection intensity (EPG) at first capture, N=88 individuals. B. Mean EPG for all individuals captured beyond first capture and after assignment to treatment categories, N=62 individuals; 166 captures C. Adult worm burden at end point for culled individuals, N=36. Data represent log means and SEM for raw EPG data. Labels above bars indicate the number of observations for each group.

**Figure S6.**
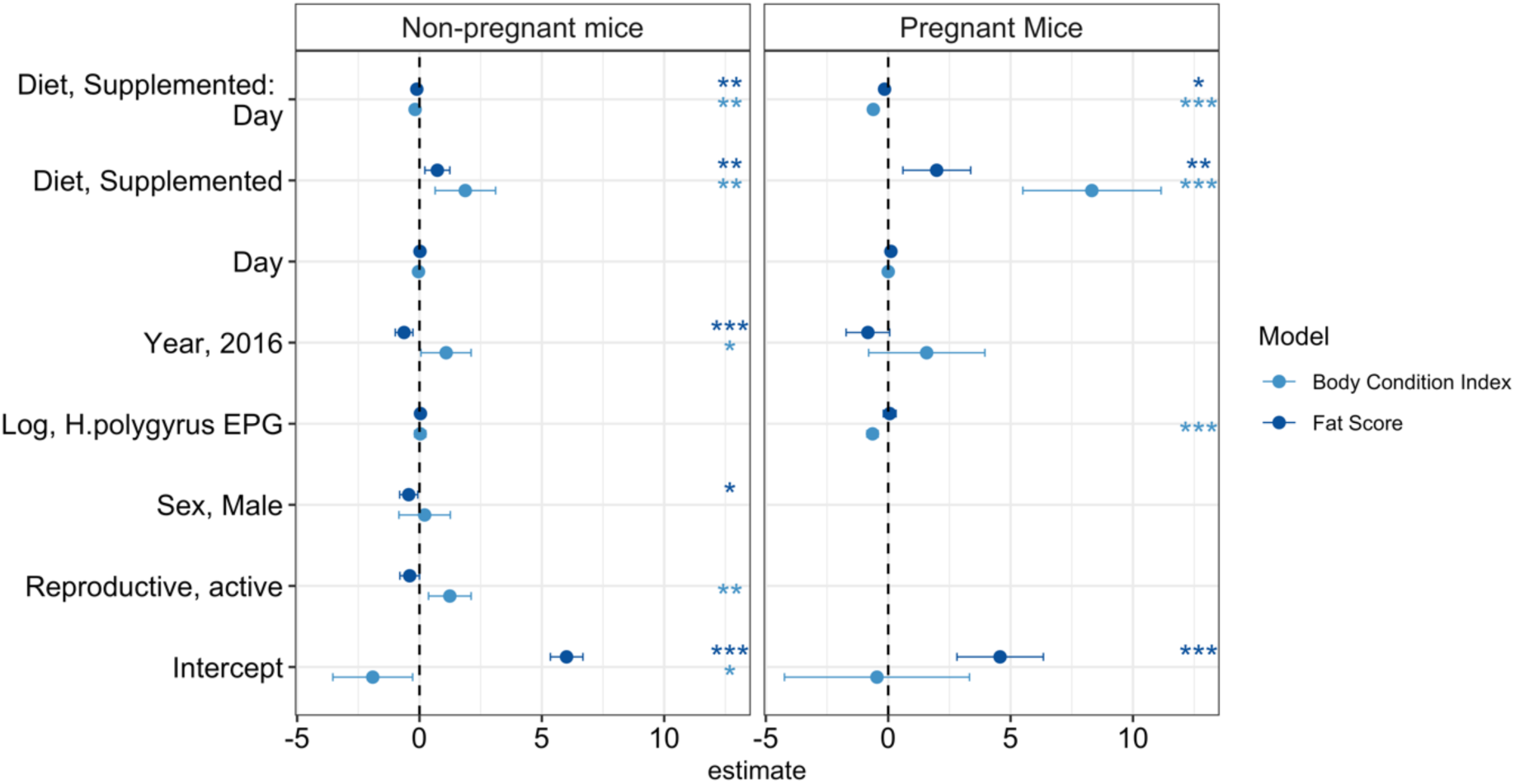
Effect size estimates from the models investigating the effect of supplemented nutrition on wood mouse body condition in the wild, measured by body condition index (residuals of weight against length regression) and total fat score. Points and ranges represent model estimates and 95% confidence intervals. Asterisks indicate the significance of variables: ***, ** and * indicate P<0.001, P<0.01 and P<0.05 respectively. Left panel represents models including all adult mice excepting pregnant females; N=79 individuals, 178 captures and right panel indicates models of pregnant females, N=15 individuals and 32 captures

**Figure S7.**
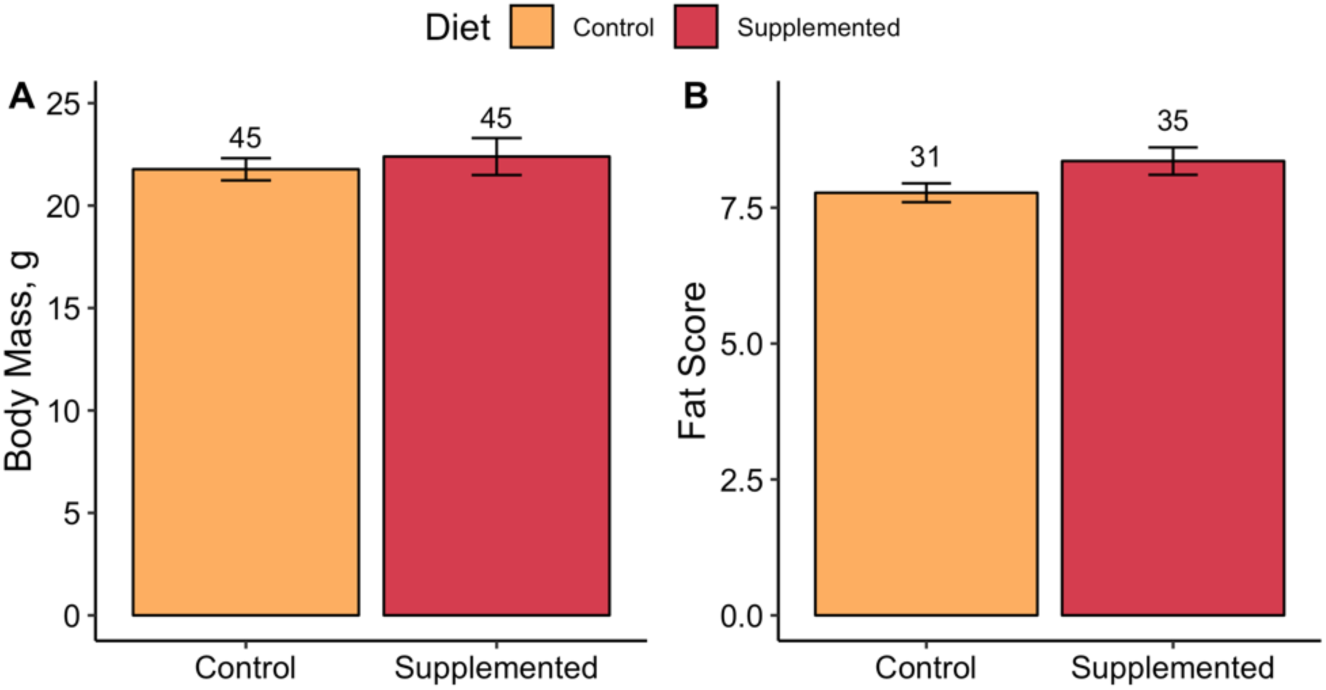
Effect of supplemented diet on body condition in laboratory wood mice. Figure represents raw means +/− SEM for (A) body mass (in g) and (B) total fat score. (2-10). Numbers above bars indicate number of observations per group.

**Figure S8.**
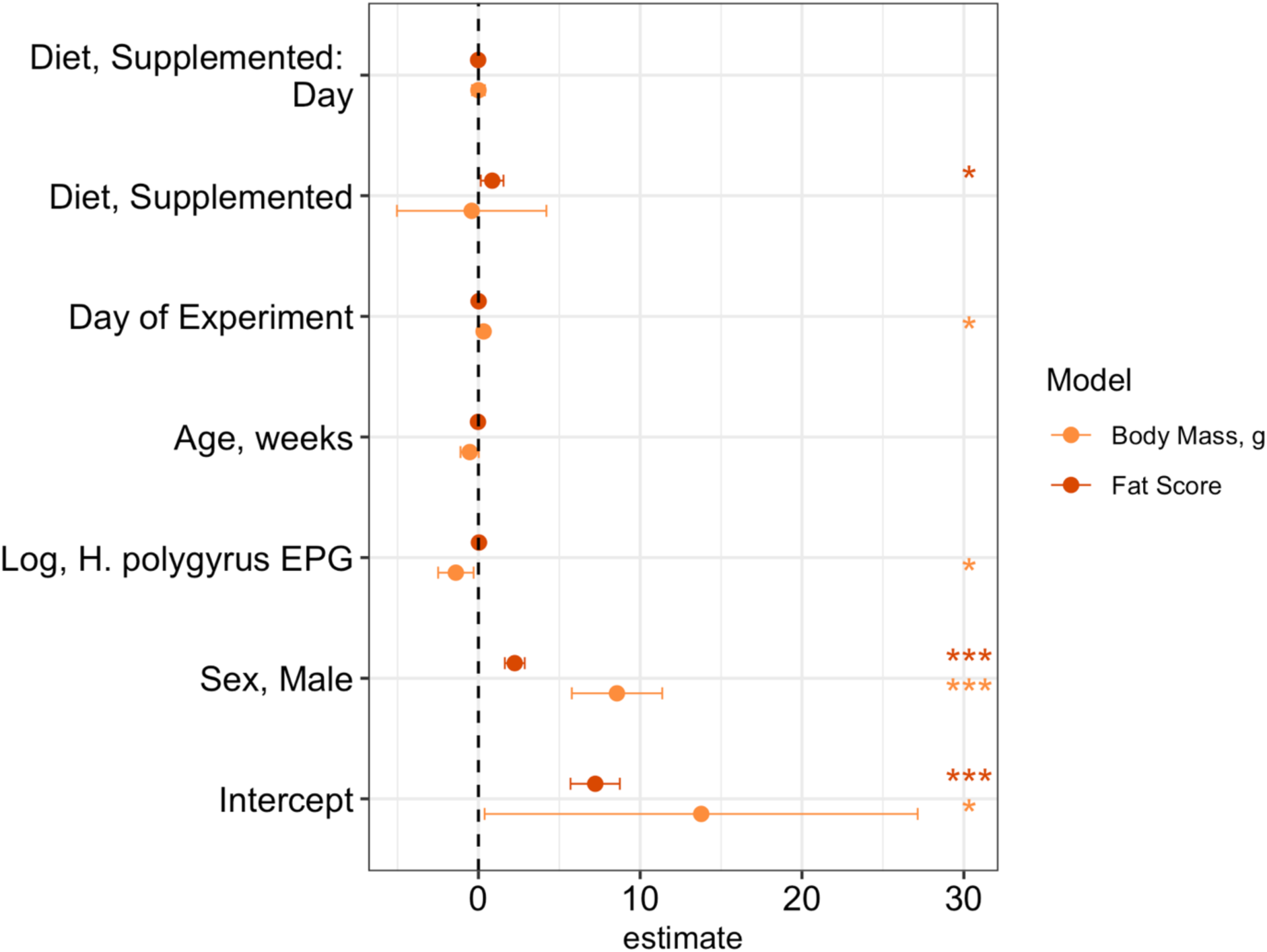
Effect size estimates from the models investigating the effect of supplemented nutrition on wood mouse body condition in the laboratory, measured by mass and total fat score. Points and ranges represent model estimates and 95% confidence intervals. Asterisks indicate the significance of variables: ***, ** and * indicate P<0.001, P<0.01 and P<0.05 respectively.

**Figure S9.**
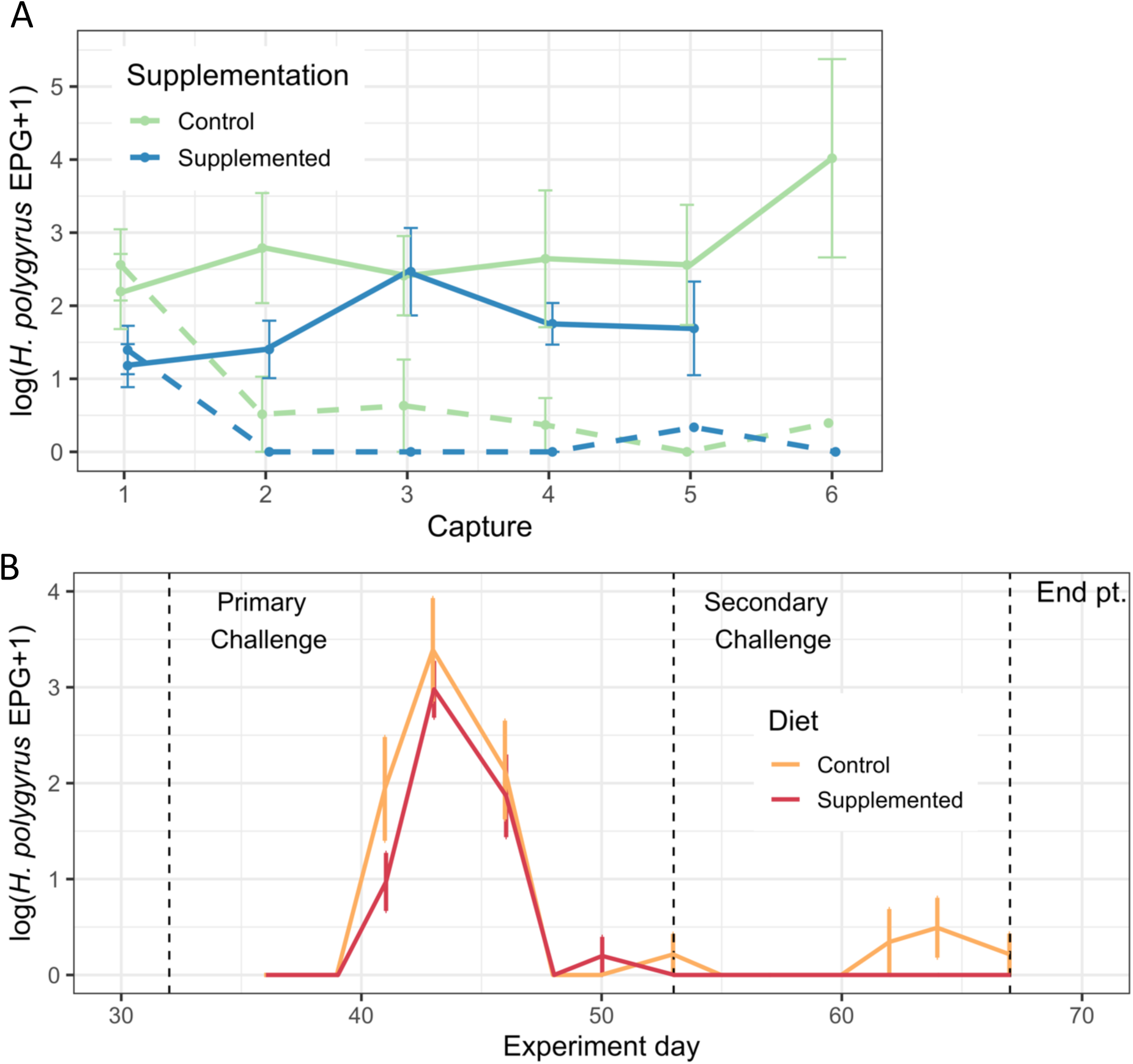
*H. polygyrus* intensity dynamics, as measured by eggs/gram over the course of (A) field experiment and (B) colony experiment. In both the wild and laboratory, supplemented mice maintain lower EPG for the majority of the experimental period. Data represent the log of EPG+1 and SE. Dashed lines in (A) represent individuals who were treated with anthelmintics. Dashed lines in (B) indicate infection and end timepoints of experiment as labelled.

